# Stabilized Full-Length Measles Fusion Protein Elicits Potent Immunity and Protection In Vivo

**DOI:** 10.1101/2025.10.07.681039

**Authors:** Dawid S. Zyla, Gillian Zipursky, Roberta Della Marca, Davide Lacarbonara, Gele Niemeyer, Weiwei Peng, Camilla Predella, Kyle Stearns, Cameron Leedale, Jonathan Miller, Daniel Tay, Jonathan Kao, Marissa Acciani, Laura Di Clemente, Anirban Das, Diptiben Parekh, Ruben Diaz Avalos, Gael McGill, N Valerio Dorrello, Kathryn M Hastie, Christopher A Alabi, Joost Snijder, Anne Moscona, Alexander L. Greninger, Stefan Niewiesk, Erica Ollmann Saphire, Matteo Porotto

## Abstract

Measles virus (MeV) is a highly contagious pathogen that causes significant morbidity and mortality in populations with low vaccination coverage. Infection typically leads to immune amnesia and, in rare cases, fatal neurological disease. While current live-attenuated vaccines are highly effective, they primarily elicit neutralizing antibodies against the hemagglutinin (H) glycoprotein, with a less robust response to the fusion (F) protein, a key protein for viral entry. To improve the immunogenicity of the F protein, we designed and characterized stabilized, prefusion MeV F protein antigens. We engineered both soluble ectodomains (F_ECTO_) and full-length, membrane-embedded proteins (F_FL_) with mutations that confer thermal stability. Cryo-electron microscopy confirmed that these engineered antigens faithfully maintain the native prefusion conformation. When evaluated in a cotton rat model, immunization with either F_ECTO_ or F_FL_ constructs induced neutralizing antibodies and elicited protection against viral challenge. The most stable full-length construct (F_FL_ 3M) elicited a more potent neutralizing antibody response than its ectodomain counterpart. Importantly, no evidence of vaccine-enhanced respiratory disease was observed. These findings establish that a thermostable, full-length F protein is a superior immunogen to its soluble ectodomain. This work presents a promising candidate for next-generation, non-replicating measles vaccines intended to complement current vaccination strategies and provide a safe option for immunocompromised individuals and others who cannot receive live-virus vaccines.

**One-Sentence Summary:** A prefusion-stabilized, full-length measles Fusion glycoprotein immunogen induces strong neutralizing responses and offers protection against challenge with wild-type virus.

## Introduction

Measles virus (MeV) is a highly transmissible morbillivirus that causes systemic disease with fever and rash. Infection of hCD150+ immune cells can deplete pre-existing memory lymphocytes, leading to “immune amnesia” and greater susceptibility to secondary infections^1^. MeV can persist in the central nervous system (CNS), causing encephalitis with long-term sequelae and, in some cases, progressing to fatal subacute sclerosing panencephalitis (SSPE). Despite long-standing availability of safe and effective live-attenuated measles vaccines, measles still causes several million cases each year and substantial mortality^2–8^.

Live-attenuated measles vaccines are given as a two-dose regimen to children older than 9 months. Many current vaccine strains are derived from the Edmonston strain, isolated in the early 1950s, and attenuated through passage. Although MeV has an RNA genome, it is genetically stable, and these vaccines continue to protect against circulating wild-type strains more than seven decades later^2–4,9,10^. Where two-dose coverage is high, regional elimination has been achieved. Outbreaks occur when coverage is insufficient. In some individuals, vaccine-induced antibody levels can wane, and revaccination with a live-attenuated vaccine often provides only modest boosting^11–17^. In addition, live vaccines are contraindicated for many immunocompromised people^4,18^. and as a result, those people who are most vulnerable to severe consequences of measles virus infection, pregnant women, pediatric cancer patients, and the immunocompromised, are the same people who cannot receive the currently available vaccine. Coupled with the very high basic reproduction number (R_0_ ≈ 12–18) and respiratory transmission, these factors support the development of complementary antigen platforms to support population immunity, especially for booster use and for those who cannot receive live vaccines.

MeV entry is mediated by two transmembrane glycoproteins on the viral envelope: hemagglutinin (H), which binds cellular receptors (hCD150 or nectin-4), and the fusion protein (F), which drives membrane fusion. F is synthesized as a precursor (F0) and cleaved by furin into a prefusion trimer of F1 and F2 subunits linked by disulfide bonds with an exposed fusion peptide at the F1 N-terminus. Prefusion F is metastable; after H engages the receptor, F refolds, exposing the fusion peptide and forming a six-helix bundle between the N-terminal (HRN) and C-terminal (HRC) heptad repeats to complete fusion^19–22^.

Neutralizing antibodies target both H and F after vaccination and natural infection. Licensed vaccines typically induce neutralization dominated by anti-H responses, while F-directed neutralization is also detectable but is more evident in naturally infected individuals who have higher overall antibody titers^23,24^. These observations suggest that presenting F in a well-defined prefusion state could broaden or strengthen humoral responses ^11–17^.

A key challenge for F-based antigens is stabilizing the prefusion state without distorting native epitopes. While several structure-guided strategies exist, some substitutions may alter antigenic surfaces^25–29^. We adopted an alternative rationale based on naturally occurring variation in clinical MeV isolates from central nervous system (CNS) infections. Sequencing of viruses from patients with measles inclusion body encephalitis (MIBE) and subacute sclerosing panencephalitis (SSPE) has repeatedly revealed hyperfusogenic substitutions in the F glycoprotein, for example L454W and T461I in the C-terminal heptad repeat (HRC), which enable cell-to-cell spread in neurons without the canonical receptors hSLAM and Nectin-4^30,31^. Additional substitutions have been identified in the HRN (*e.g.,* G168R, E170G) and at the protomer interface (*e.g.,* S262G) ^32^. While hyperfusogenic changes promote neuronal spread, they often destabilize F and decrease viral stability ^33^. We also identified complementary mutations that stabilize the prefusion state, such as E170G, which was found in an SSPE case ^32^. Moreover, during the rescue of recombinant viruses carrying the L454W mutation, a compensatory substitution, E455G, emerged. Mutations such as E170G and E455G counterbalance the destabilizing effects of hyperfusogenic mutations. For example, the combination of L454W and E455G produces an F protein with thermal stability comparable to wild-type F ^33^.

Building on our prior identification of the two stabilizing substitutions described above (E170G and E455G)^32–35^, we engineered prefusion-stabilized ectodomains (F_ECTO_) and full-length, membrane-embedded F (F_FL_), and in some constructs, we also removed the furin cleavage site to prevent F0 processing.

We produced and purified both F_ECTO_ and F_FL_ in the prefusion state, confirmed conformation by cryo-electron microscopy, and quantified stability by biochemical differential scanning fluorimetry. We also profiled N-glycosylation and modeled potential glycan shielding. Abolishing the cleavage site markedly increased stability, with the largest effect in the full-length protein.

In cotton rats, immunization with stabilized F_ECTO_ or F_FL_ elicited neutralizing antibodies and reduced virus replication after challenge. The most stabilized full-length construct induced higher neutralizing titers than the matched ectodomain. These findings identify a thermostable, prefusion full-length MeV F as a strong antigen candidate for future vaccine development.

## Results

### Age-dependent neutralization titers correlate with F-specific antibodies

The measles-mumps-rubella (MMR) vaccine is highly effective, primarily through induction of neutralizing antibodies. The precise targets and function of these antibodies against currently circulating wild-type viruses are still being defined, including their roles in blocking receptor engagement and membrane fusion. Prior studies indicate that most vaccine-elicited neutralizing activity is directed to the hemagglutinin (H) glycoprotein, with variable contributions from antibodies to the fusion (F) protein^23,24^. In contrast, F-directed neutralization is present in natural infection^23^. Further, overall measles antibody titers are also typically higher after infection than after vaccination^23^. As the proportion of donors with natural infection declines, pooled human sera for post-exposure prophylaxis may show lower neutralization potency^4,36^. We set out to understand these differences.

To understand in detail which components of the polyclonal human immune response are linked to measles virus neutralization, and for whom, we profiled humoral responses in 51 MeV-seropositive individuals (ages 1–76 years, 18 females and 33 males). We measured serum binding to vaccine virus (Moraten) H and F proteins, each as encoded by vaccine virus (Moraten), wild-type virus (B3, one of two currently circulating strains), and neuropathogenic virus (B3 F L454W)^30,33,34^ strains. For each human serum sample, we also measured neutralization titers (IC_50_) against the wild-type B3 virus, as well as inhibition of Measles-driven cell–cell fusion. The strongest correlation with neutralizing circulating wild-type B3 virus is serum binding to the vaccine-strain H protein (Spearman ρ=0.571, p=1.2×10^−5^; n=51; Fig. 1A, Fig. S1, Table S1). Binding to wild-type F protein shows a weaker, but significant association with neutralization (ρ = 0.381; p = 0.006), whereas binding to vaccine F shows only a trend (ρ = 0.265; p = 0.06; Fig. 1B-C, Table S1). Binding to the postfusion–biased mutant F L454W does not correlate with neutralization (ρ=-0.049; p=0.731; Fig. 1D).

**Figure 1.**
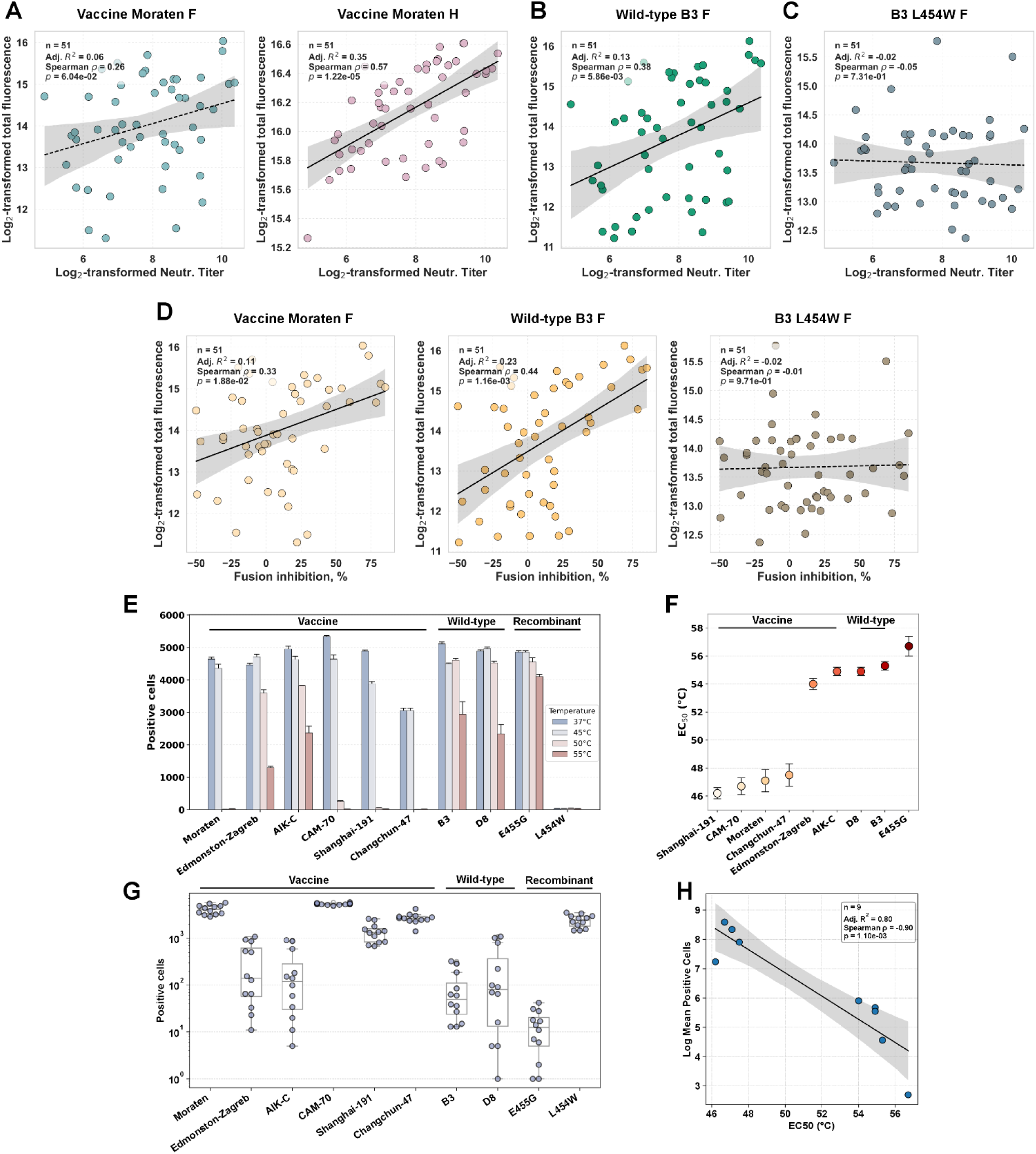
Human serology and heat-challenge assays relating antibody binding, neutralization, and prefusion F stability. (A) Correlations between serum binding to vaccine-strain measles glycoproteins and neutralization titers. Points show log₂-transformed integrated fluorescence for binding to Moraten vaccine F (left) or Moraten vaccine H (right) plotted against log₂-transformed neutralization titers (neutralization titer; IC₅₀ of reciprocal serum dilution log₂(neutralization titer+1)). Lines indicate least-squares fits with 95% confidence bands; per-panel R², Spearman ρ, and p values are shown. (B) Binding to wild-type B3 F versus log₂ (neutralization titer+1). (C) Binding to destabilized L454W wild-type B3 F versus log₂ (neutralization titer+1). (D) Binding to F versus inhibition of cell–cell fusion in an H head–independent assay. Left, vaccine F (Moraten); center, wild-type B3 F; and right, destabilized L454W wild-type B3 F. The fusion readout uses a chimeric attachment protein (MeV H stalk/NDV head) that is not recognized by anti-MeV H head antibodies; negative values indicate more fusion than the no-serum control. Lines show least-squares fits with 95% confidence bands; R², ρ, and p values are reported on each plot. (E) Cell-surface heat-stability of F proteins from vaccine strains (Moraten, Edmonston-Zagreb, AIK-C, CAM-70, Shanghai-191, Changchun-47), wild-type strains (B3, D8), and controls (stabilized E455G; destabilized L454W). Transfected cells were incubated for 30 min at the indicated temperatures (37, 45, 50, and 55 °C) and stained with prefusion-specific mAb 77; bars represent the number of mAb 77-positive cells. Data represent three independent experiments; error bars denote standard error. (F) The stability of the tested fusion glycoproteins was assessed by their EC_50_ values, determined through fitting a 4-parameter logistic function to data from (E). This model shared the same top (A), bottom (B), and Hill slope (p) across constructs, with only the inflection temperature (x₀, representing EC₅₀) varying. The destabilized L454W construct was excluded from the fitting process because it lacked a detectable prefusion state, even at the lowest temperature tested. Error bars indicate the standard error of the measurements. (G) Cell-surface staining for post-triggered state of F proteins from vaccine strains (Moraten, Edmonston-Zagreb, AIK-C, CAM-70, Shanghai-191, Changchun-47), wild-type strains (B3, D8), and controls (stabilized E455G; destabilized L454W). Transfected cells were incubated overnight at 37°C and stained with post-triggered mAb C6; bars show the number of mAb C6-positive cells in a logarithmic scale. Data includes all points from three independent experiments with all 3 replicates; the line inside the box denotes the median, the box bounds indicate the interquartile range (IQR; 25th–75th percentiles), and the whiskers extend to the most extreme values within 1.5× IQR (H) Relationship between prefusion heat stability and accumulation of post-triggered F on the cell surface. The x-axis shows EC₅₀ (°C) extracted from panel F; the y-axis shows the log_2_ mean number of C6 (post-triggered-specific) mAb-positive cells from panel G. Each point represents one F variant (n = 9; L454W excluded because EC₅₀ could not be defined). The line denotes a least squares fit with the 95% confidence band. Higher EC₅₀ (greater prefusion stability) correlated with lower C6 binding (adjusted R² = 0.80; Spearman ρ = −0.90, p = 1.1 × 10⁻³).

To distinguish F-directed from H-directed neutralization, we used a fusion assay with a chimeric attachment protein containing the MeV H stalk and the Newcastle disease virus (NDV) globular head. The NDV head is not recognized or neutralized by antibodies directed against the measles H head region^37^. In this H-head-independent context, serum binding to wild-type F correlates with fusion inhibition (Spearman ρ=0.442, 95% CI 0.172-0.659; p=0.001) and remains stronger than vaccine F (ρ=0.328, 95% CI 0.057-0.565; p=0.019; **Fig. 1C**, **Fig. S1**, **Table S1**).

The age of the serum sample donor matters, with different responses observed at different ages, and notably, older individuals exhibit the greatest F-mediated neutralization. Using log-transformed neutralization titers (log2(IC_50_+1)), age-stratified regressions revealed distinct correlation patterns between antigen-specific binding and neutralization (**Fig. S2, S3**). For sera of young people, aged 1-25 years, neutralization correlates strongly with vaccine H binding (ρ = 0.67, Adj. R² = 0.55, p = 2.0 × 10⁻³). There is no significant association of neutralization with binding of vaccine F in young subjects (Adj. R² = 0.003, p = 0.33) or B3 F (Adj. R² = 0.09, p = 0.17). For adults 26-40 years old, none of the antigens, including vaccine F, vaccine H, and wild-type B3 F, showed statistically significant associations (all Adj. R² ≤ 0.04, p > 0.25). Middle-aged adults 41-53 years old exhibited the strongest correlations between F-specific binding and neutralization: with binding of wild-type B3 F the strongest correlate (Adj. R² = 0.56, p = 7.8×10⁻³), followed closely by binding of vaccine F (Adj. R² = 0.52, p = 0.012), followed by binding of H (Adj. R² = 0.44, p = 0.026). In older adults (ages 56–76), binding to vaccine (Moraten) F, vaccine (Moraten) H, and wild-type (B3) F each correlates significantly with neutralization (vaccine Moraten F: adjusted R² = 0.50, ρ = 0.745, p = 0.013; vaccine Moraten H: adjusted R² = 0.41, ρ = 0.690, p = 0.027; B3 F: adjusted R² = 0.415, ρ = 0.693, p = 0.026, **Table S1**). Binding to the postfusion L454W B3 mutant F (neuropathogenic variant) does not correlate with neutralization in older adults (adjusted R² = 0.165, ρ = 0.508, p = 0.134) or any other age stratum (all p ≥ 0.13; adjusted R² ≤ 0.17), individually or all ages combined (all ages combined: adj. R² = −0.019, p = 0.813).

For anti-F antibodies, a key mechanism of action may be blockage of F-mediated membrane fusion. Hence, we next analyzed the correlation of antigen binding to fusion inhibition. Binding of B3 wild-type F is the most predictive of fusion inhibition in middle-aged and older adults: ages 41–55 (Adj. R² = 0.543, p = 2.0×10⁻³) and ages 56–76 (Adj. R² = 0.742, p = 8.37×10⁻⁴). For the 56–76-year-old cohort, vaccine (Moraten) F binding also exhibits a strong association (Adj. R² = 0.725, p = 1.0 × 10⁻³), with vaccine (Moraten H) binding being additionally significant but less predictive (Adj. R² = 0.438, p = 2.2 × 10⁻²).

Together, these data indicate that neutralization in younger donors, who are primarily vaccine-primed, is largely H-associated, with little measurable dependence on F. In contrast, in older adults (who might have been clinically or subclinically infected or, if vaccinated, unknowingly boosted by wild-type virus), neutralization is more associated with binding of F. For these older adults, binding of wild-type F is more predictive of neutralization than binding of vaccine F, and is always more predictive than binding of F in the postfusion conformation. These observations support presenting F in a stabilized, prefusion form as a potential booster antigen.

### Thermal stability of prefusion F detected by prefusion-specific antibody

To explore the differences between attenuated vaccine and wild-type F, we next asked whether F proteins from vaccine *vs.* wild-type strains differ in thermal stability and presentation of the prefusion conformation. In these experiments, we transfected cells with two strains of wild-type measles (B3 and D8) and 6 types of vaccine (Moraten, CAM-70, Shanghai-191, Changchun-47, Edmonston-Zagreb, AIK-C), or with two control versions of F, stabilized in the prefusion conformation (E455G) or the postfusion conformation (L454W). To determine the stability of the different forms of F expressed on the cell surface, we assessed the prefusion state under heat stress using a conformation-specific monoclonal antibody (mAb 77) that binds a prefusion-specific epitope, which is lost in the conformational change to the postfusion state. Twenty-four hours after transfection and incubation at 37 °C, cells expressing vaccine-strain Fs, circulating wild-type F, or control versions of F were exposed at 45, 50, or 55 °C for 30’ and then stained with mAb 77 to assess the prefusion-positive cells (**Fig. 1E-F, Fig. S4**).

At 37-45 °C, most versions of F, across various strains, exhibit robust binding of mAb 77. At 50 °C, F from several vaccine strains (Moraten, CAM-70, Shanghai-191, Changchun-47) lose binding to mAb 77, consistent with the loss of the prefusion-specific epitope, and suggesting refolding to the postfusion state. In contrast, the circulating wild-type B3 and D8 F proteins and vaccine strains AIK-C and Edmonston-Zagreb (albeit at lower levels) retain mAb 77-positive cells at 50°C. At 55 °C, only the stabilized E455G control F still shows strong mAb 77 reactivity (∼80%). The destabilized L454W control F was essentially non-reactive at all tested temperatures.

These data indicate that, under our expression and assay conditions, F from both wild-type measles virus strains and one vaccine strain preserve consistent prefusion epitopes at elevated temperatures. Most vaccine strains lose prefusion presentation at elevated temperature. The engineered stabilized control performs best, and the destabilized control performs worst. These data support the view that wild-type F constructs are expressed in a prefusion state that is more stable than most of the vaccine strains, including the reference vaccine strain (*i.e.,* Moraten). Based on these findings, we asked whether the vaccine F is presented in the postfusion state at 37 °C. To test whether vaccine Fs display post-trigger features at 37 °C, we probed cell-surface F with mAb C6, which recognizes a post-triggered epitope^38^ (**Fig. 1G**). C6 bound to all the F proteins, but separated into three tiers: (i) high reactivity to C6 for Moraten, CAM-70, Shanghai-191, Changchun-47, similar to the destabilized L454W control; (ii) low binding to C6 for AIK-C and Edmonston-Zagreb, similar to wild-type B3 and D8; and (iii) minimal binding to C6 for the stabilized E455G control. C6 reactivity at 37 °C inversely tracked thermal retention of the prefusion-specific mAb 77 epitope (ρ = −0.90, adjusted R² = 0.80, p = 1.1×10⁻³), indicating that greater prefusion stability accompanies reduced exposure of the post-trigger epitope (**Fig. 1H**).

Together, these findings suggest that several vaccine F proteins adopt the post-triggered state at physiological temperature and further support stabilized prefusion F as a booster antigen candidate.

### Structural and biophysical analysis of purified and stabilized F ectodomains

We previously published a stabilized variant of wild-type F (IC323 strain, Uniprot: Q786F3) containing two mutations, E170G and E455G, termed F_ECTO_ 2M^35^. This F includes natural substitutions instead of the disulfide bridges present in the stalk region previously tested^39^. These natural E to G substitutions stabilize the prefusion state while still permitting conformational transition to postfusion, albeit in a more gradual and controllable manner than wild-type F^35^. Here, F_ECTO_ 2M was modified by eliminating the furin cleavage site to generate F_ECTO_ 3M (F_ECTO_ 2M + Δfur, **Fig. 2A–B**), to reduce the aggregation propensity previously observed during refolding of F_ECTO_ 2M. F_ECTO_ 2M and 3M were expressed in Drosophila S2 and ExpiCHO cells, respectively, and purified for biophysical analysis. Thermal stability and aggregation were assessed using nano-differential scanning fluorimetry (nanoDSF) and dynamic light scattering (DLS). Removal of the furin cleavage site increased the apparent melting temperature from 49.2 °C (F_ECTO_ 2M) to 51.9 °C (F_ECTO_ 3M) (**Fig. 2C**, top). DLS further showed that F_ECTO_ 3M maintained a stable cumulant radius of ∼30 nm up to 80 °C, whereas F_ECTO_ 2M aggregated above 50 °C (**Fig. 2C**, bottom). These results indicate that removing the furin cleavage site increases thermal stability and reduces aggregation, yielding a better-behaved antigen for storage, routine handling, and downstream characterization, including immunization.

**Figure 2.**
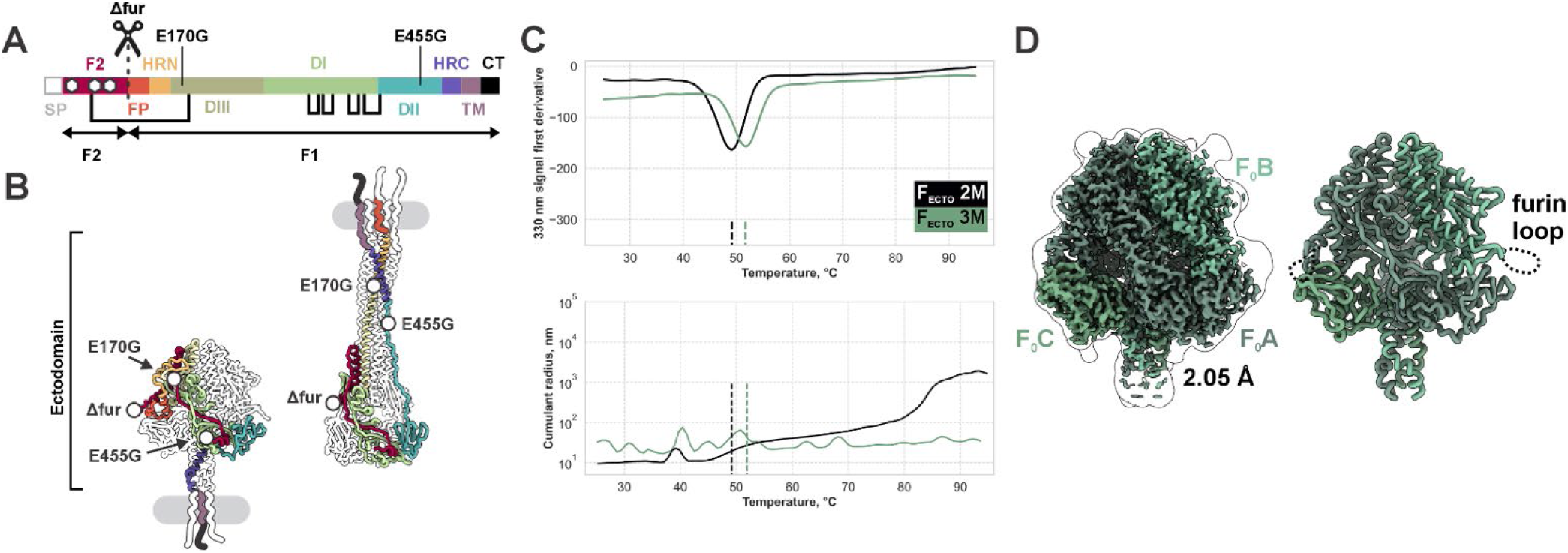
Biophysical Characterization and Structure of Stabilized Measles F Antigens. (A) Domain map of full-length F (F_FL_). Color-coded schematic showing SP (signal peptide), F2 (with N-glycans marked as white hexagons), FP (fusion peptide), HRN, DI–DIII, HRC, TM, and CT. Native disulfides (black links) and the furin cleavage site that generates F2/F1 are indicated. Stabilizing changes are labeled: Δfur (cleavage-site deletion), E170G, and E455G. (B) Location of stabilizing substitutions. Prefusion ectodomain models with sites E170G, E455G (2M), and Δfur (3M) highlighted. The gray bars depict the membrane plane; unmodeled segments are shown schematically. Coloring matches panel A. (C) Thermal and colloidal behavior of purified ectodomains (F_ECTO_). Top, nanoDSF first-derivative traces for F_ECTO_ 2M (black) and F_ECTO_ 3M (green). Apparent melting temperatures (mean of three experiments) are 49.2 °C (2M) and 51.9 °C (3M). Bottom, DLS cumulant radius versus temperature. F_ECTO_ 2M shows a radius increase consistent with aggregation above ∼50 °C, whereas F_ECTO_ 3M remains ∼30 nm up to ∼90 °C. Dashed vertical lines mark the nanoDSF T_m_ values from the top panel. (D) Cryo-EM structure of F_ECTO_ 3M. Prefusion trimer resolved to 2.05 Å. Left, cryo-EM map colored by protomer (F₀A–C) with a Gaussian-blurred silhouette. Right, corresponding ribbon model; dashed loops denote unresolved segments that include the mutated furin-cleavage region.

To further assess the impact of the furin cleavage site on protein structure, we solved the cryo-EM structure of F_ECTO_ 3M to 2.05 Å resolution. The structure of F_ECTO_ 3M, missing the cleavage site, is highly similar to that of the previously determined F_ECTO_ 2M (global RMSD ∼0.3 Å), which has an intact cleavage site (**Fig. 2D, Fig. S5A, Fig. S6A**). Minor structural differences were localized to a flexible loop (residues 164-172) and the furin cleavage site region itself, consistent with previously observed variations in F_ECTO_ 2M (**Fig. S6**). Hence, deletion of the furin cleavage motif does not affect the majority of F structure or F antibody epitopes^35,38,40^.

### Characterization of full-length stabilized F constructs

We purified membrane-embedded full-length F (F_FL_) variants and compared them to their ectodomains (**Fig. 3A– D, Fig. S8**). Thermal stability was measured by nanoDSF and DLS (**Fig. 3A**). F_FL_ 2M showed a slightly higher apparent melting temperature than its ectodomain counterpart (50.2 °C *vs*. 49.2 °C), whereas F_FL_ 3M was marginally lower than F_ECTO_ 3M (51.6 °C *vs*. 51.9 °C). DLS trends were consistent with nanoDSF, indicating comparable colloidal stability across full-length and ectodomain formats in this temperature range.

**Figure 3.**
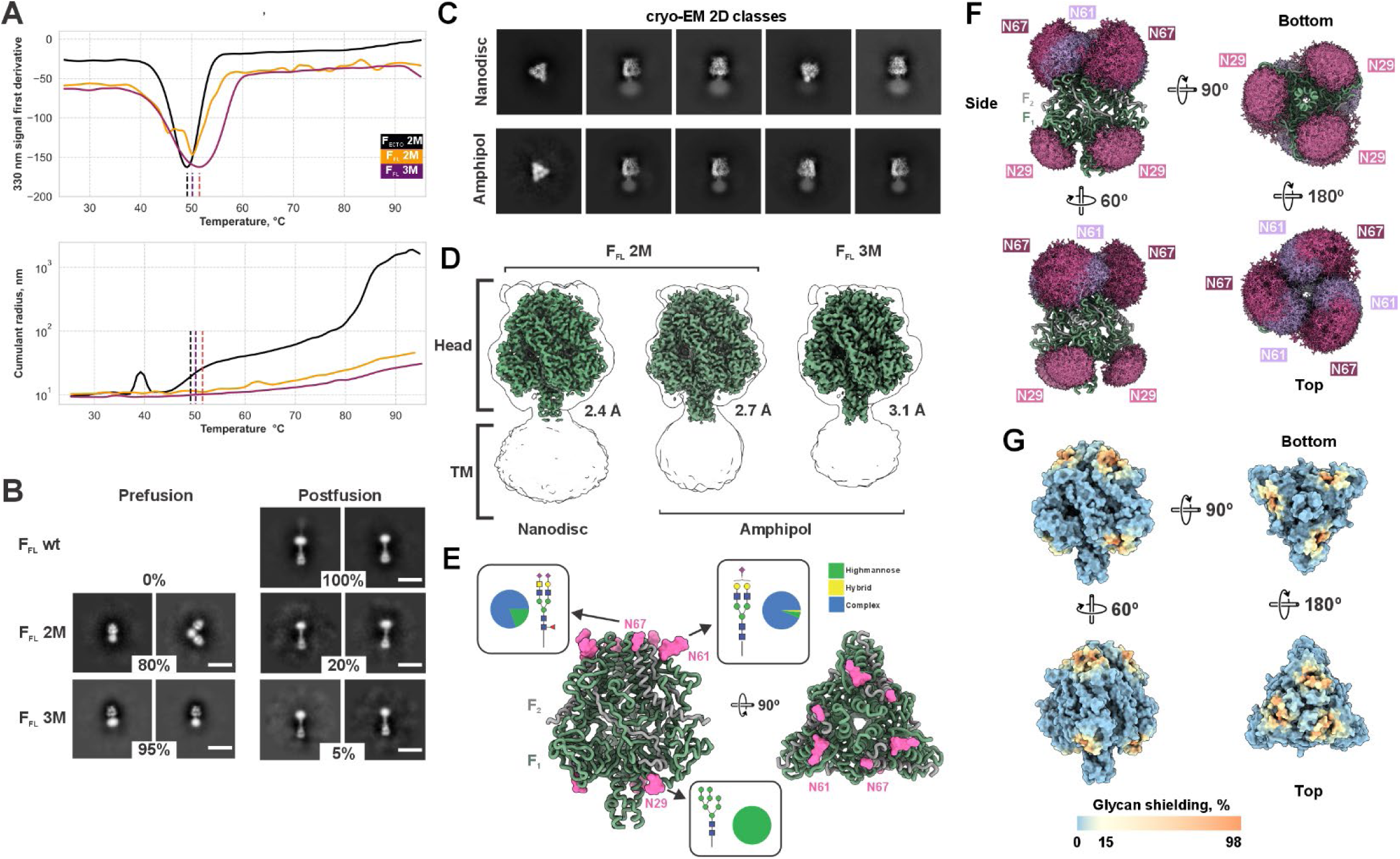
Structural characterization of full-length measles F (F_FL_) and glycan mapping. (A) Thermal and colloidal behavior. Top, nanoDSF first-derivative traces for F_ECTO_ 2M (ectodomain; black), F_FL_ 2M (magenta), and F_FL_ 3M (orange). Apparent melting temperatures (mean of three experiments): 49.2 °C (F_ECTO_ 2M), 50.2 °C (F_FL_ 2M), 51.6 °C (F_FL_ 3M). Bottom, DLS cumulant radius versus temperature. F_ECTO_ 2M shows a radius increase consistent with aggregation above ∼50 °C, whereas both full-length constructs remain comparatively stable across the range shown. Dashed lines mark the nanoDSF T_m_ values. (B) Prefusion/postfusion populations assessed by negative-stain EM. Representative 2D classes and estimated proportions for F_FL_ wild-type (0% prefusion), F_FL_ 2M (80% prefusion / 20% postfusion), and F_FL_ 3M (95% / 5%). Classes were derived from ∼100 micrographs per construct with similar initial particle counts. Scale bar, 200 Å. (C) Cryo-EM 2D classes of F_FL_ 2M in nanodiscs (top) and amphipol (bottom). Head features are well defined; TM-region detail is limited. (D) Reconstructions of full-length F. Head-only high-resolution maps for F_FL_ 2M in nanodiscs (2.4 Å) and amphipol (2.7 Å), and for F_FL_ 3M in amphipol (3.1 Å). A Gaussian-blurred silhouette (1.5 σ, gray outline) depicts low-resolution density attributable to the membrane region/TM, which is not resolved. (E) Glycosylation of the F_FL_ head (2M). Ribbon model colored F1 (green) and F2 (gray) with three N-linked sites per protomer highlighted (magenta). Pie charts summarize site-specific compositions from MS: green, high mannose; yellow, hybrid; blue, complex. Monosaccharide symbols: blue square, GlcNAc; red triangle, fucose; green circle, mannose; yellow circle, galactose; yellow square, GalNAc; purple diamond, Neu5Ac. (F) Glycan envelope. Conformer libraries were placed at N29 (basal), N61, and N67 (apical) using GlycoSHIELD. N61/N67 form a dense apex barrier; N29 is predominantly high-mannose and is located near the base. (G) Modeled glycan shielding. Surface representation colored by estimated shielding percentage (scale: blue, minimal; yellow, low; orange, high), highlighting exposed and glycan-occluded regions across multiple orientations.

Negative-stain EM quantified conformational populations after membrane extraction (**Fig. 3B**). Wild-type full-length F was exclusively in the postfusion state. In contrast, 80% of F_FL_ 2M and 95% of F_FL_ 3M proteins were prefusion, confirming the stabilizing effect of the two mutations in the context of the full-length protein and indicating that Δfur further enhances structural stability.

For high-resolution analysis, F_FL_ was reconstituted in nanodiscs or amphipol (**Fig. 3C–D**). Cryo-EM resolved the head and stalk domains but not the transmembrane (TM) region. To improve TM visualization, we employed computational approaches (including focused refinements and flexibility-targeted pipelines) as well as experimental strategies (addition of the prefusion-stabilizing peptide [FIP–HRC]₂–PEG_11_ ^41^ and replacement of nanodiscs with amphipol), as detailed in **Fig. S5B–E**, **S6B–E**, and **S8A–B**. Despite these efforts, the TM region remained unresolved, suggesting substantial structural heterogeneity and/or the absence of a defined quaternary arrangement. Amphipol reconstructions of F_FL_ 2M and F_FL_ 3M revealed partial density proximal to the membrane (residues 488–491; **Fig. S8B–D**). The backbone deviates from the trajectory of residues 484–486, consistent with a bend and a helix discontinuity between the stalk and the TM helix, a feature not resolved in prior ectodomain maps. Bioinformatic analyses indicate that similar membrane-proximal flexibility may occur in other paramyxovirus F proteins (**Fig. S8E–F**). Across constructs, the prefusion head architecture of full-length F closely matched that of the ectodomains (**Fig. 3D, Fig. S7B–D**), with no detectable alterations attributable to stabilizing mutations. Thus, stabilization enhances the prefusion state without changing the presentation of head epitopes relevant for antibody recognition.

### Glycan Composition and Epitope Shielding of MeV F

We used mass spectrometry to characterize the glycosylation patterns of mammalian cell-expressed F_FL_ 2M (**Fig. 3E**) to explore the impact of glycosylation on epitope availability, as glycosylation patterns and their shielding effects influence both antigen stability and epitope accessibility^42–45^. We found that MeV F has an estimated 8 kDa of glycans per protomer, resulting in a ∼12% glycan mass fraction. With only three N-linked glycosylation sites per protomer (N29, N61, and N67), the glycan mass fraction is relatively low compared to other viral glycoproteins such as HIV gp160 (42%), Lassa GP (33%), Mumps (22%), or Nipah (20%). The glycans linked to N29 of MeV F are exclusively high mannose, possibly due to obstructed access to the site after protein folding. In contrast, those linked to N61 and N67 are predominantly complex in nature. While the majority of glycans at N67 bear a single (core) fucose, a core fucose is largely absent from N61. Both glycans at N61 and N67 are predominantly di-antennary structures, carrying one or two terminal N-Acetylneuraminic acid residues. While the antennae at N61 are predominantly galactosylated, at N67 we observe a large portion of glycans bearing N-Acetylgalactosamine, forming a LacdiNAc structure in one of the arms instead. Modeling the fully glycosylated MeV F using the single most abundant glycoforms at each of the three sites reveals that the glycans linked to N61 and N67 shield most of the apex of F, while the rest of the glycoprotein’s face is exposed for immune recognition (**Fig. 3F-G**).

### Stabilized F_FL_ constructs elicit both *in vivo* protection and neutralizing titers

Prior studies with RSV demonstrate that recombinant stabilized F ectodomains can protect after vaccination^27,29,46–50^. Based on these results, we investigated whether stabilized MeV F elicits neutralizing antibodies and whether a soluble ectodomain (F_ECTO_) or a full-length, membrane-embedded antigen (F_FL_) is a more effective immunogen.

Cotton rats were immunized with F_ECTO_ 2M, F_ECTO_ 3M, F_FL_ 2M, or F_FL_ 3M (5 µg per dose, two doses). A licensed live-attenuated comparator was included (Edmonston-Zagreb). Animals were challenged with wild-type B3-eGFP Measles virus (**Fig. 4**). F_ECTO_ 2M, F_ECTO_ 3M, and F_FL_ 3M each conferred complete protection, comparable to the live-attenuated Edmonston-Zagreb comparator (**Fig. 4A**). F_FL_ 2M yielded partial protection (2/5 fully protected). After dose two, neutralizing titers were significantly above mock for F_ECTO_ 2M, F_ECTO_ 3M, and F_FL_ 3M. Further, after dose two, F_FL_ 3M produced the highest neutralizing titers, exceeding those of the matched F_ECTO_ 3M ectodomain and the full-length Edmonston-Zagreb vaccine comparator group (≈1:1,000 *vs.* ≈1:180; **Fig. 4B**). Note that the comparator is a complete virus, and its neutralization titer reflects combined anti-H plus anti-F responses, while responses from F immunization alone are anti-F alone, and lack a neutralization contribution from H.

**Figure 4.**
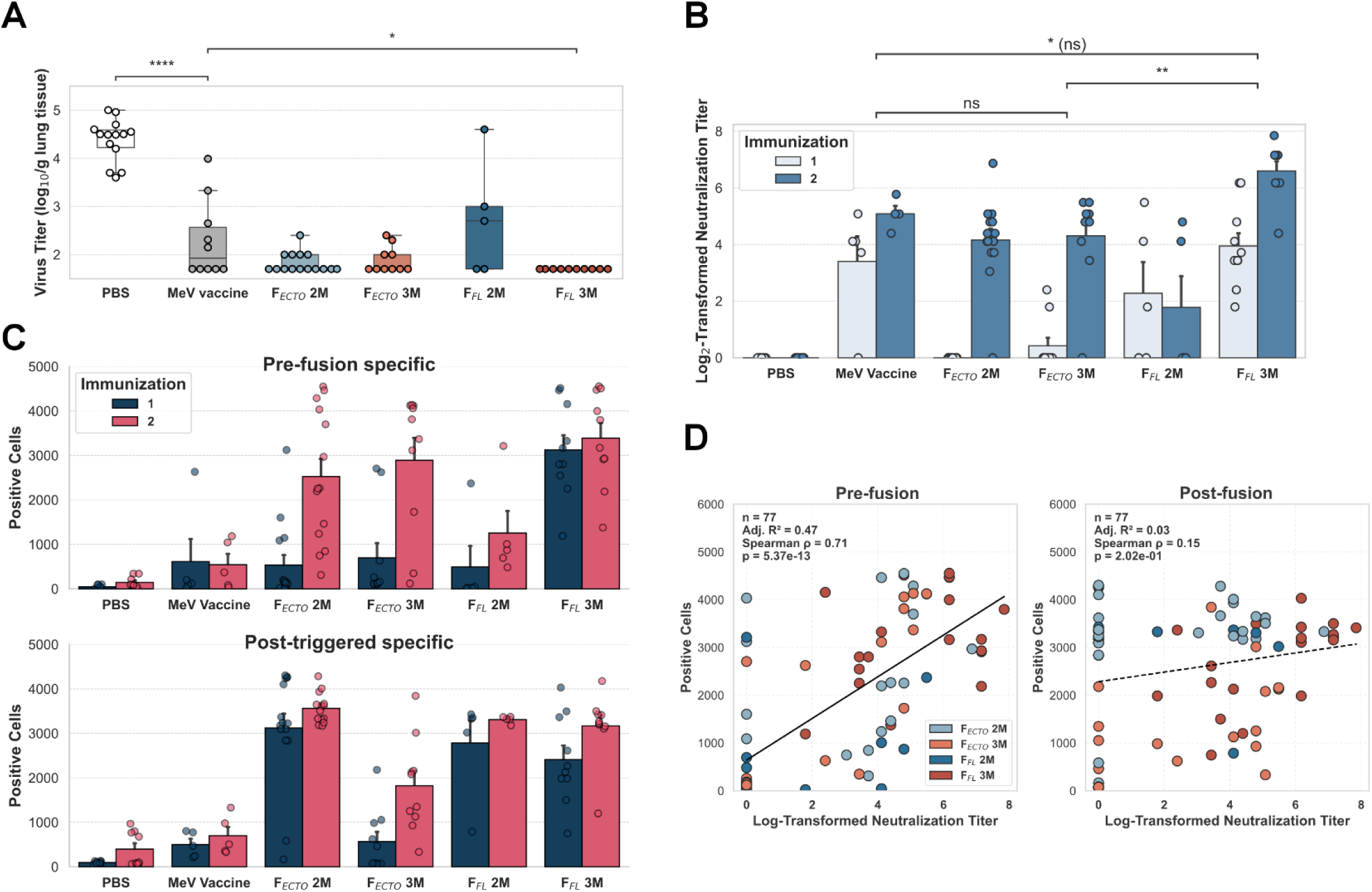
Cotton rat immunization with stabilized measles F antigens: protection, neutralization, and epitope specificity. (A) Challenge outcome. Cotton rats received two doses (4-week interval) of 5 µg protein in alum—F_ECTO_ 2M, F_ECTO_ 3M, F_FL_ 2M, or F_FL_ 3M, or 10⁴ TCID₅₀ of the Edmonston-Zagreb (EZ) vaccine; PBS served as control. Animals were challenged intranasally with wild-type B3-eGFP measles virus. Lung virus titers are shown with a detection limit of 2 log₁₀ PFU/g; values below the line indicate undetectable titers. Statistics for the second immunization only used two-sided Mann–Whitney–Wilcoxon tests with Holm– Bonferroni correction (ns (p_adj_ > 0.05), * (0.01 < p_adj_ ≤ 0.05), ** (0.001 < p_adj_ ≤ 0.01), *** (0.0001 < p_adj_ ≤ 0.001), **** (p_adj_ ≤ 0.0001). (B) Neutralizing titers. Log₂-transformed neutralization titers (IC₅₀ of reciprocal serum dilution) four weeks after each dose. Points denote individual animals; bars, means; error bars, 95% CI. After two doses, F_FL_ 3M achieved the highest titers among protein groups; the difference *vs.* F_ECTO_ 3M was significant after correction (**). Differences *vs.* the EZ comparator did not reach significance after correction. (C) Conformation-specific binding. Sera from PBS, EZ, and protein-immunized animals were tested for binding to cells expressing wild-type F. Prefusion epitopes were measured at 37 °C; post-triggered epitopes were induced by heating to 55 °C for 30 min. Bars show means with standard error for the first (dark) and second (light) immunizations. Prefusion recognition increased most for F_ECTO_ 3M and F_FL_ 3M after the boost; post-triggered recognition was higher for the less stabilized 2M constructs. (D) Link between prefusion recognition and neutralization. Correlations between log₂ neutralization titer (from B) and conformation-specific binding (from C) for protein groups (PBS and EZ excluded). Prefusion binding correlated with neutralization (R²=0.47, p=5.05×10⁻¹²; solid fit), whereas postfusion binding did not (R²=0.04, p=7.68×10⁻²; dashed fit). Shaded bands indicate 95% confidence intervals.

Sera from the most stable immunogens (F_ECTO_ 3M, F_FL_ 3M) preferentially recognize prefusion F, even after a single dose, whereas F_FL_ 2M sera preferentially recognize post-triggered F (**Fig. 4C**). The full-length presentation of the most stabilized F (F_FL_ 3M) further increased neutralization relative to its matched ectodomain (F_ECTO_ 3M). Prefusion recognition correlates with stronger neutralization (**Fig. 4D**), indicating that stabilizing F in the prefusion state enhances functional antibody responses.

To assess the risk of vaccine-enhanced disease^51^, we examined lung histopathology after challenge (**Fig. S9**). Animals immunized with F subunit proteins (whether F_ECTO_ or F_FL_) showed no evidence of atypical measles or enhanced pathology. Peribronchiolar infiltrates were similar to the live-attenuated group, and substantial perivasculitis or interstitial pneumonitis was not observed (**Table S3**).

## Discussion

Live-attenuated measles vaccines remain the foundation of measles control and have dramatically reduced global disease burden. In settings where vaccination coverage cannot be maintained, however, population-level immunity can decline, and outbreaks may follow upon introduction by travelers. CDC data show that measles vaccination coverage among kindergartners was above 95% before 2019 and has declined to below 93% in the 2024–2025 period. A recent model predicts that measles could become endemic in the U.S. if current low vaccination rates persist^52^. Infants too young to be vaccinated and individuals who cannot receive live vaccines remain particularly vulnerable^2,4,18^. These realities require alternate complementary antigen approaches that can bolster population immunity without replacing licensed vaccines.

Here, we developed a highly immunogenic F protein stabilized in the ideal conformation to elicit neutralizing and protective antibodies. We define substitutions that stabilize measles F in both ectodomain and full-length constructs, validate their thermal and structural properties, and demonstrate *in vivo* activity as recombinant protein antigens in cotton rats. The most stable full-length construct (F_FL_ 3M) elicited potent neutralizing antibodies. It provided complete protection from the B3 wild-type virus challenge, with neutralizing titers that exceeded those induced by the vaccine strain vaccinated group in this model. Because only F was administered in these groups, the measured neutralization and protection reflect F-directed immunity alone. Protection from F-mediated immunity alone is notable, as vaccine-mediated neutralization was previously attributed largely to H, with F thought to contribute little or to a much lower extent. To be considered a vaccine candidate, the prefusion F protein must be further evaluated in non-human primates. Furthermore, the prefusion state is preferable to the postfusion state, as indicated by the different levels of neutralizing antibodies elicited by the F_FL_ 3M compared to the F_FL_ 2M. This is consistent with previous vaccine development efforts, particularly for RSV, that demonstrated considerable protection with a recombinant ectodomain of the fusion protein^28,53^, especially when stabilized in its prefusion conformation^54–56^. Our findings also highlight an important finding for measles: the full-length, membrane-embedded F outperformed the stabilized ectodomain. This suggests that the membrane-proximal region, together with the continuous display of prefusion-sensitive epitopes in a native-like context, may be crucial for optimal immune recognition. Importantly, Δfur stabilization enhanced thermal tolerance and increased prefusion state without disrupting the conformation of the prefusion head, supporting the preservation of key antigenic sites. Furthermore, cell-based heat-challenge assays showed that while wild-type F can retain prefusion epitopes at elevated temperatures, engineered stabilization confers the most durable presentation. Together, these results suggest that for measles vaccine strategies, prioritizing the native full-length F protein may yield superior immunogenicity. Importantly, there is a fundamental distinction between RSV and MeV: RSV F alone mediates viral entry, whereas MeV F is responsible for fusion, with attachment mediated by the H protein. Consequently, a subunit-based measles vaccine would likely require inclusion of both H and F. However, our data indicate that antibodies targeting F alone are sufficient to block infection.

The analysis of human sera in these experiments provided additional context. In younger, vaccine-primed donors, neutralization was strongly associated with binding to the vaccine H protein and showed little correlation with binding to the F protein. In contrast, in older adults, who may have endured measles or been unknowingly and subclinically exposed to circulating measles, binding to F became predictive of neutralization. It remains to be determined whether the reduced thermal stability of most vaccine F proteins, relative to wild-type strains, contributes to the skewed H-based neutralization observed in younger individuals, and overall lower neutralizing titers; a possibility that will require further investigation. Low measles seropositivity has been observed in children despite two or more doses of live-attenuated vaccine, with minimal improvement from additional doses, highlighting gaps in our understanding of suboptimal vaccine responses^12^. These findings underscore the potential of presenting F in a stabilized prefusion conformation to enhance F-mediated neutralization, suggesting that optimizing F stability could be key to broadening and strengthening protective immunity.

In summary, our data identify a thermostable, full-length prefusion MeV F as a strong antigenic candidate intended to complement licensed vaccines. The work introduces a stabilization strategy grounded in naturally occurring compensatory substitutions, provides high-resolution structural validation, and demonstrates *in vivo* immunogenicity and protection without evidence of enhanced respiratory disease. The advances presented here support further development of stabilized F, possibly in combination with recombinant H^57^, in future non-replicating booster formats and for populations unable to receive live vaccines, or in boosters for those who did not respond adequately to live-attenuated vaccines.

## Acknowledgments

The authors thank NIAID R56AI183536 and R01AI190902 (MP and EOS), NIAID R01AI176833 and U19AI181984 (MP), NINDS R01NS105699 and R01NS091263 (MP), NIAID R21AI180456 (EOS), Swiss National Science Foundation Postdoc Mobility fellowships P2EZP3_195680 (DSZ) and P500PB_210992 (DSZ), and Institutional Funds of the La Jolla Institute for Immunology (EOS) for funding, and Dr. Ruben Diaz and the LJI Cryo-EM Core for data collection. Instrumentation in the cryo-EM core was supported by U19 AI109762-S1, gifts from the GHR Foundation, and private philanthropic support. JS and WP are supported by the Dutch Research Council NWO Gravitation 2013 BOO, Institute for Chemical Immunology ICI 024.002.009. Biospecimens and/or data utilized for this research were obtained from the Columbia University Biobank (CUB), which is partially supported by Columbia University’s Clinical and Translational Science Award (CTSA) funded through Grant Number UL1TR001873.

We thank Nanotemper for access to Prometheus Panta. We thank D. Deneka and C. Alvadia for their suggestions regarding the nanodisc assembly protocol. We thank F.T. Bovier for technical assistance. We greatly appreciated Rik de Swart’s constructive feedback on the manuscript. We are also grateful to M. Mor for carefully reading the manuscript and to H. M. Callaway and H. Li for helping at the early stages of the project.

## Supplementary figures

**Figure S1.**
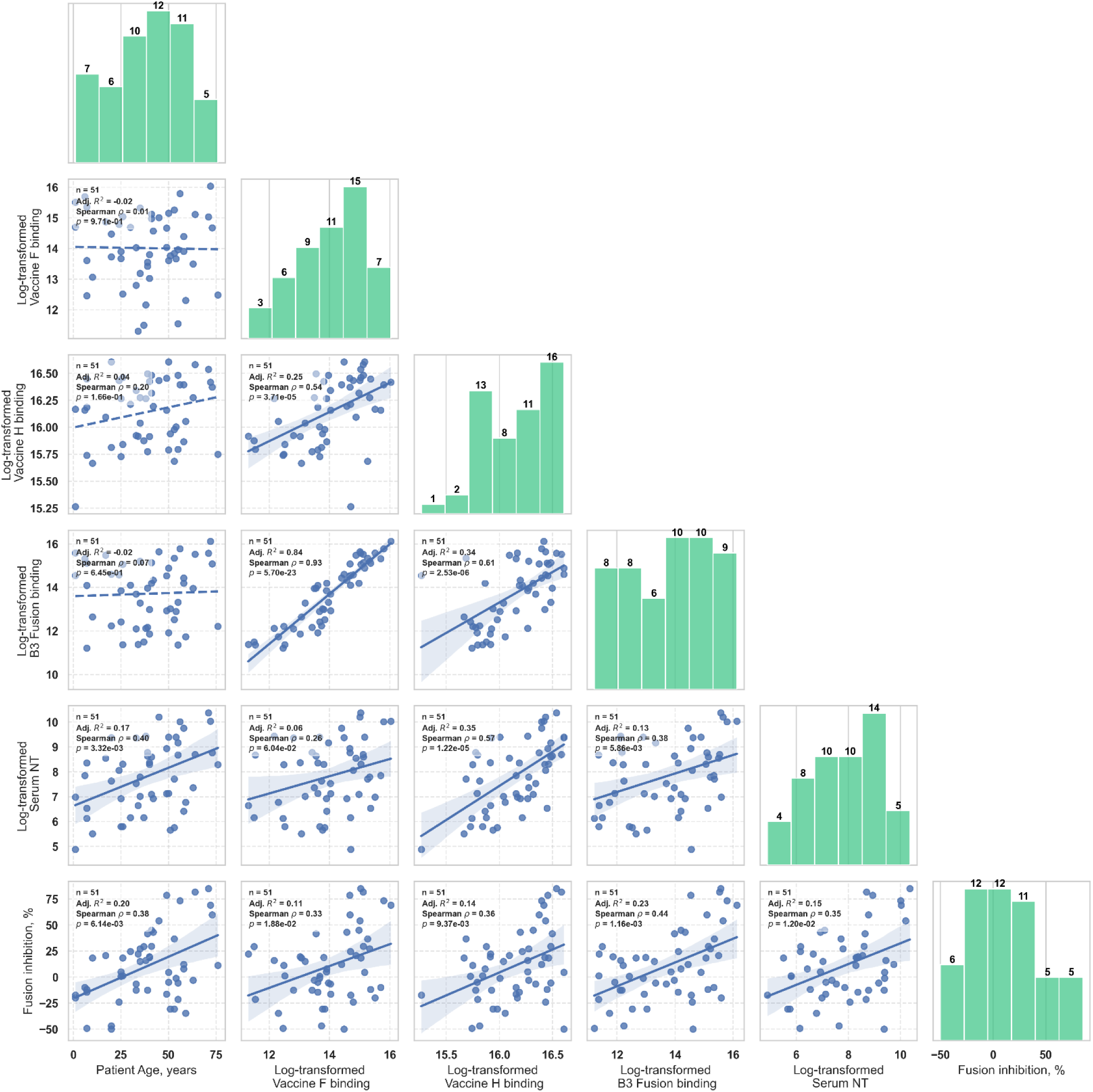
Pair plot illustrating the relationships between all measured assays and patient age. The diagonal displays the distribution of each variable. Significant correlations are represented by solid lines with a light blue overlay, indicating the 95% confidence interval of the regression estimate. Non-significant correlations are depicted with dashed lines. Histogram plots on the diagonal include bar labels that represent the values within each bin.

**Figure S2.**
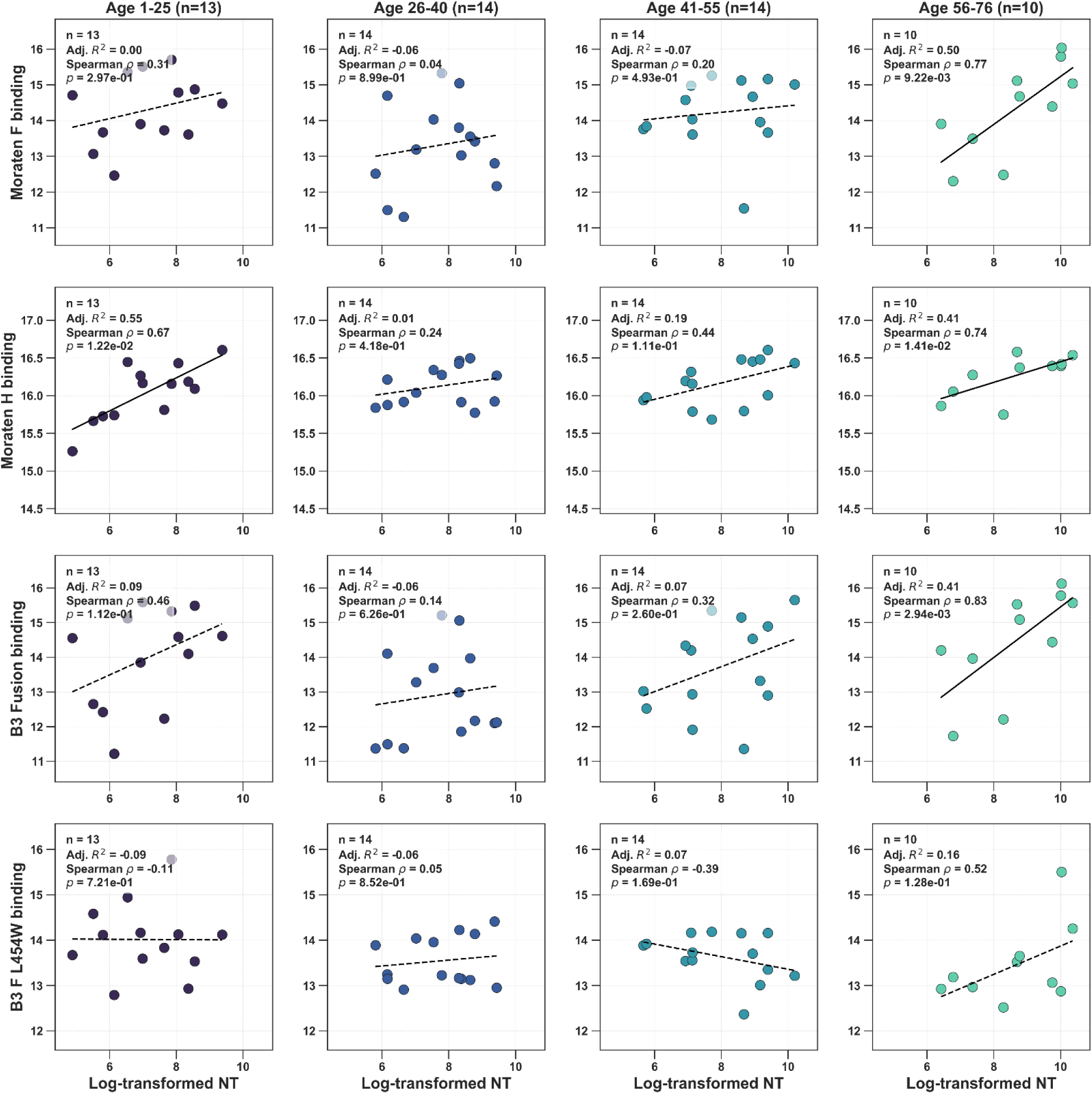
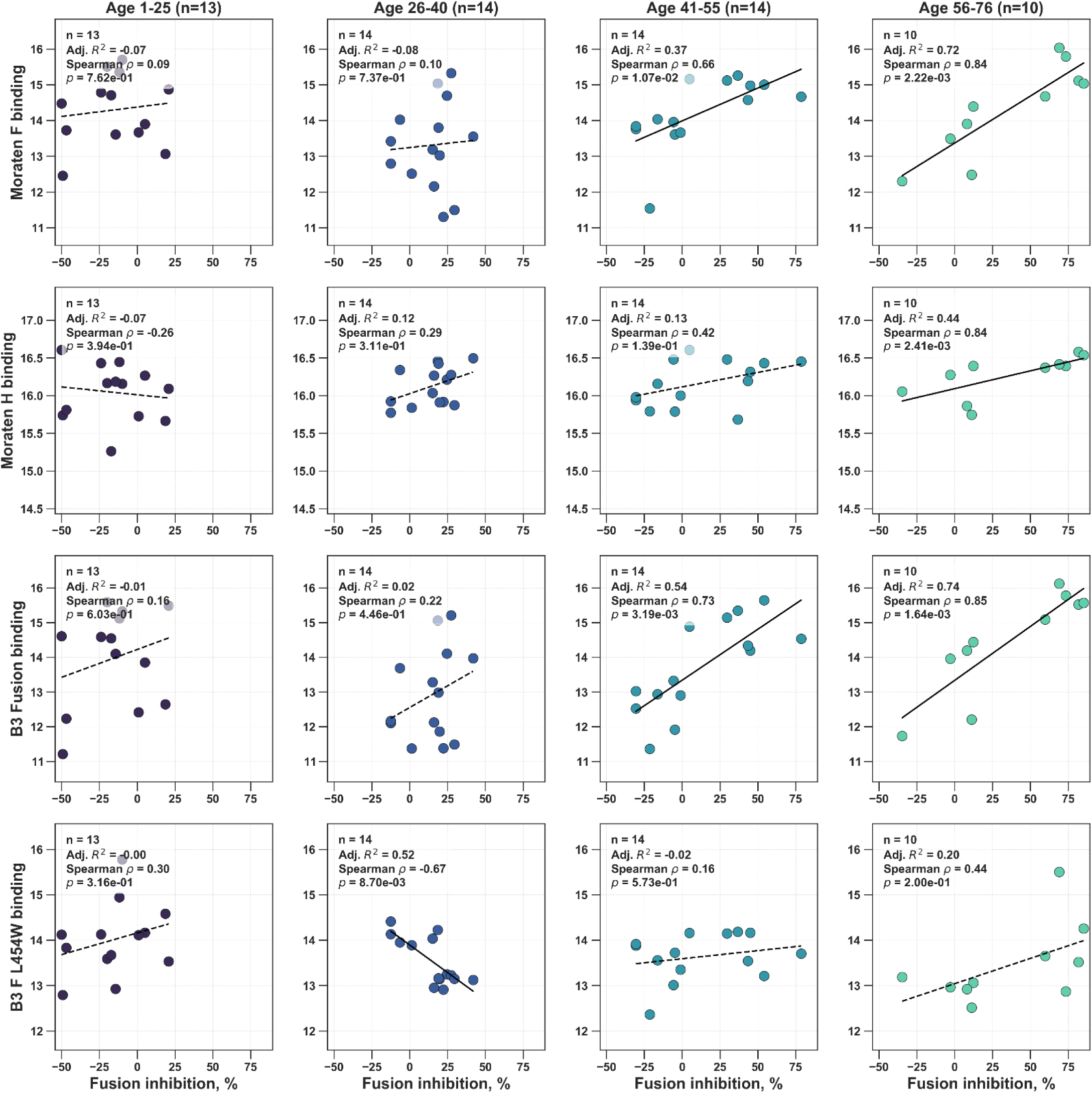
Age-stratified correlations between antigen binding, neutralization, and fusion inhibition. Scatter plots by age group—1–25 (n=13), 26–40 (n=14), 41–53 (n=14), and 54–90 (n=10)—showing log₂-transformed binding (integrated fluorescence) versus log₂-transformed neutralization titers (NT, upper) or fusion inhibition (%, lower). Rows correspond to binding targets: vaccine F (top), vaccine H (second from the top), wild-type B3 F (second from the bottom), and destabilized B3 L454W F (bottom). Lines denote least-squares fits; per-panel R², ρ, and p values from linear regression are reported. Axes are scaled consistently within each row and column.

**Figure S3.**
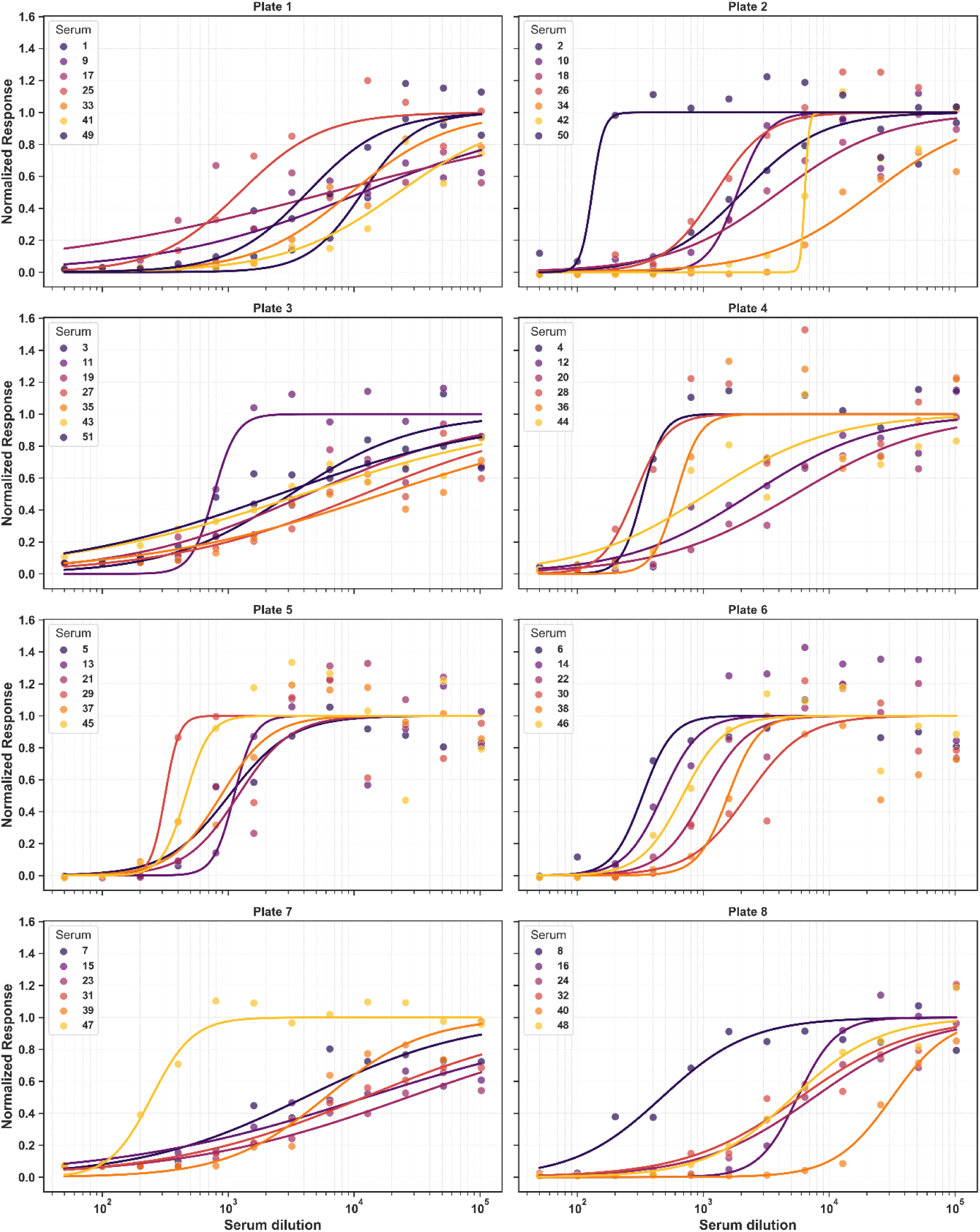
Fitting of serum viral neutralization assay data. A logistic function with globally shared minimum and maximum signal parameters was applied to fit the data for each serum dilution per plate. All values were normalized using the parameters derived from the fit to ensure consistent data representation.

**Figure S4.**
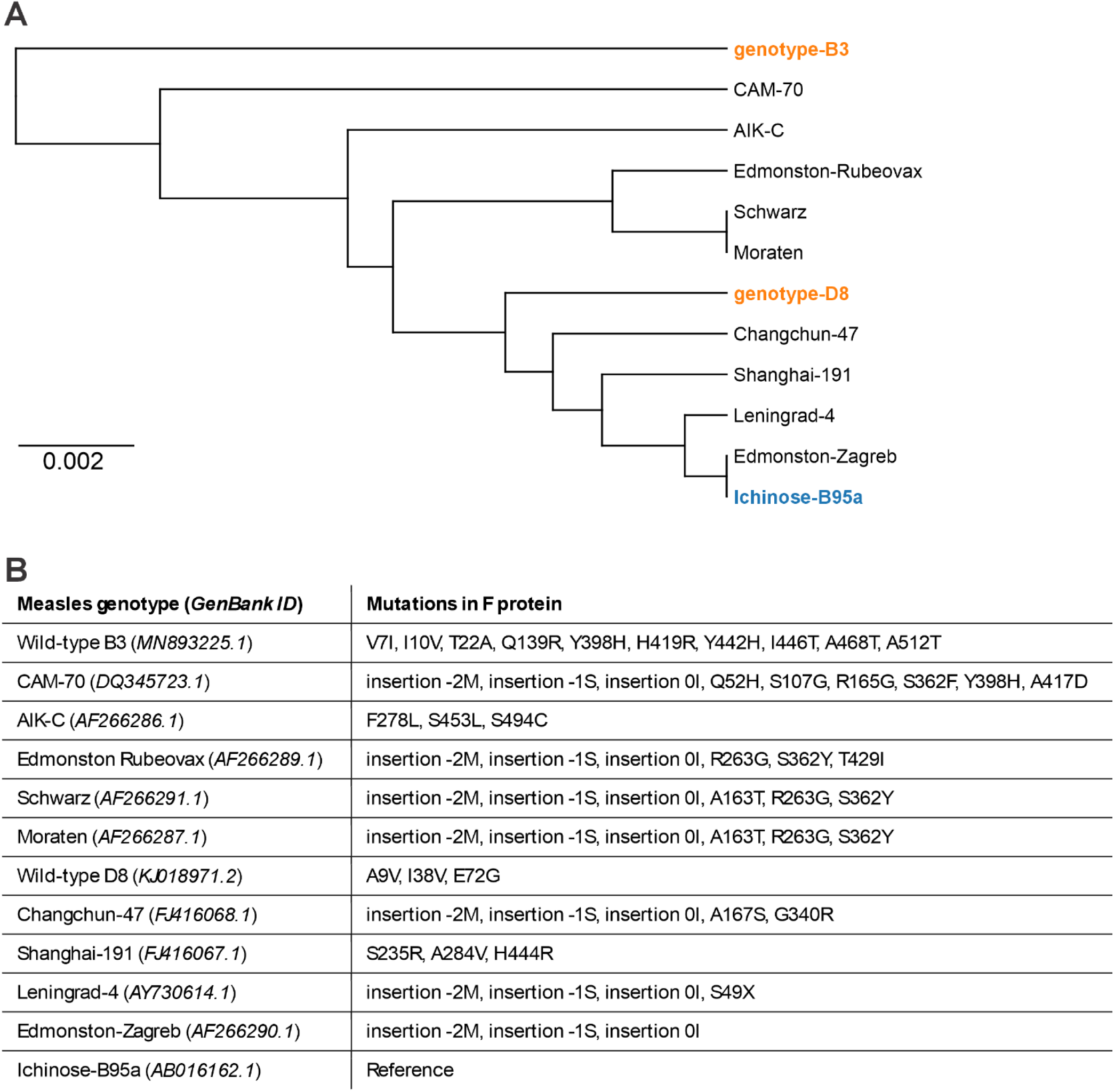
Phylogeny and naturally occurring variation in the measles F protein. (A) Unrooted phylogenetic tree derived from a multiple-sequence alignment of representative measles F proteins. The reference sequence (Ichinose-B95a) is highlighted in blue; circulating wild-type genotypes are highlighted in orange (*e.g.,* B3, D8). Vaccine-lineage strains (*e.g.,* Schwarz, Moraten, Edmonston variants, CAM-70, AIK-C) are shown in black. Scale bar indicates substitutions per site (0.002). (B) Amino-acid differences in F relative to the Ichinose-B95a reference with GenBank IDs shown. Listed changes include single-residue substitutions and N-terminal insertions; the notations “insertion −2M/−1S/0I” denote additional residues upstream of the reference N terminus as annotated in the source sequences. “X” indicates an undetermined residue in the deposited sequence.

**Figure S5.**
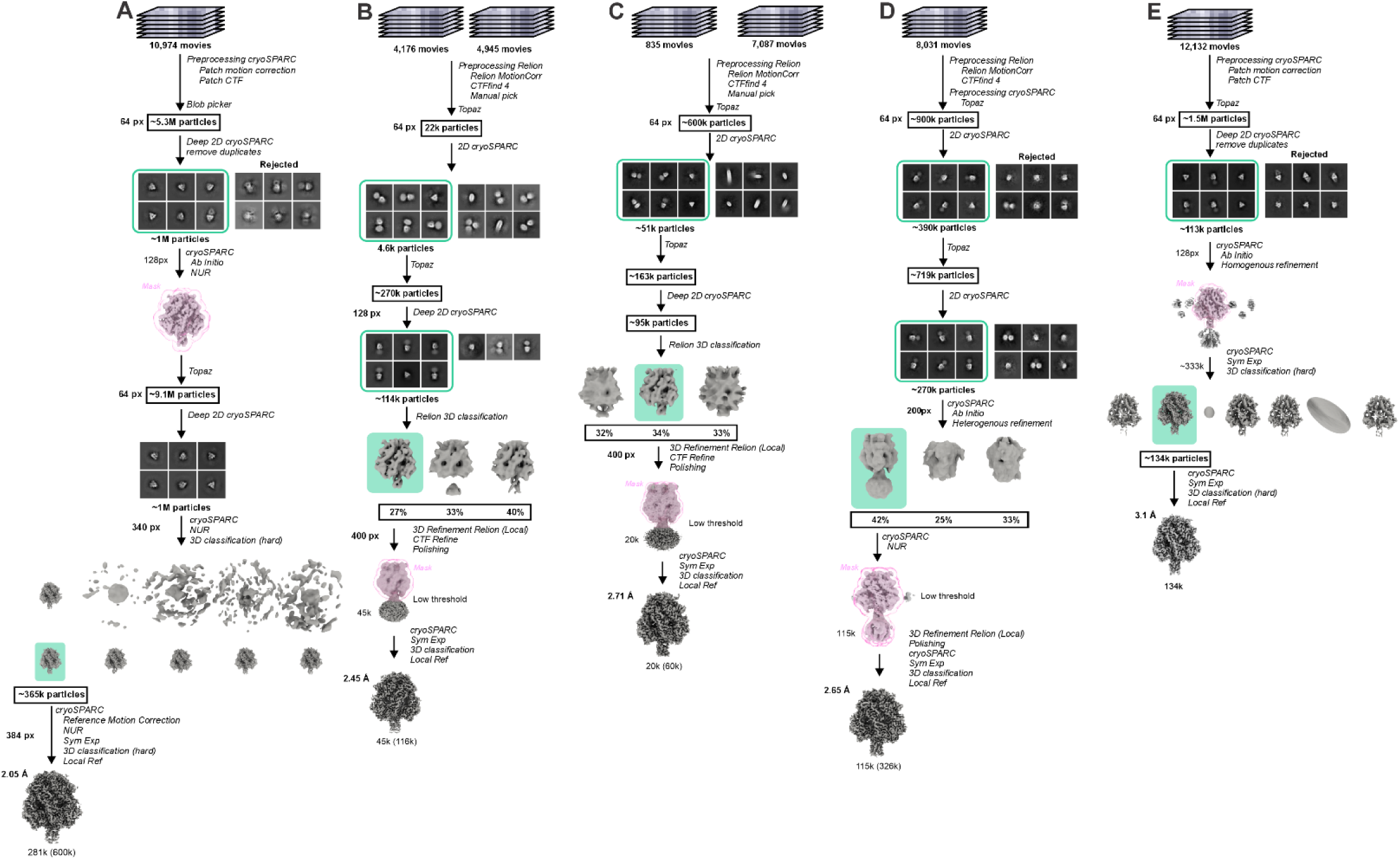
Cryo-EM data processing flowchart. Selected classes are shown in green. A) F_ECTO_ 3M B) F_FL_ 2M in nanodisc C) F_FL_ 2M + [FIP-HRC]_2_-PEG_11_ in nanodisc D) F_FL_ 2M + [FIP-HRC]_2_-PEG_11_ in amphipol E) F_FL_ 3M in amphipol

**Figure S6.**
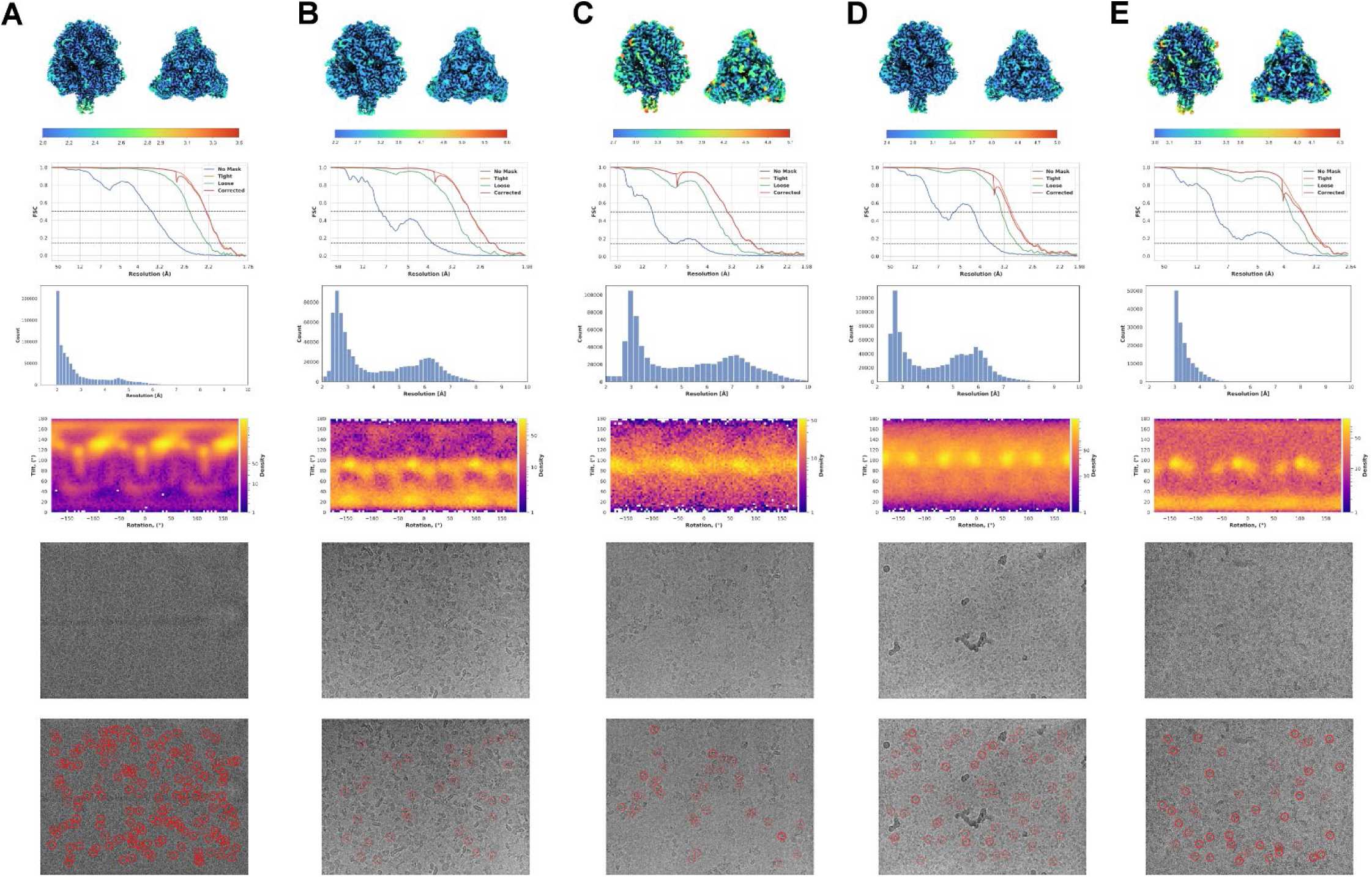
Cryo-EM statistics of recorded datasets. The first row corresponds to local resolution mapped on the final volume, the second shows the Fourier shell correlation, the third shows local resolution histogram derived from the data in row 1, the fourth row shows the angular distribution, and the fifth and sixth rows show representative micrograph and particle locations present in the final particle stack. A) F_ECTO_ 3M B) F_FL_ 2M in nanodisc C) F_FL_ 2M + [FIP-HRC]_2_-PEG_11_ in nanodisc D) F_FL_ 2M + [FIP-HRC]_2_-PEG_11_ in amphipol E) F_FL_ 3M in amphipol

**Figure S7.**
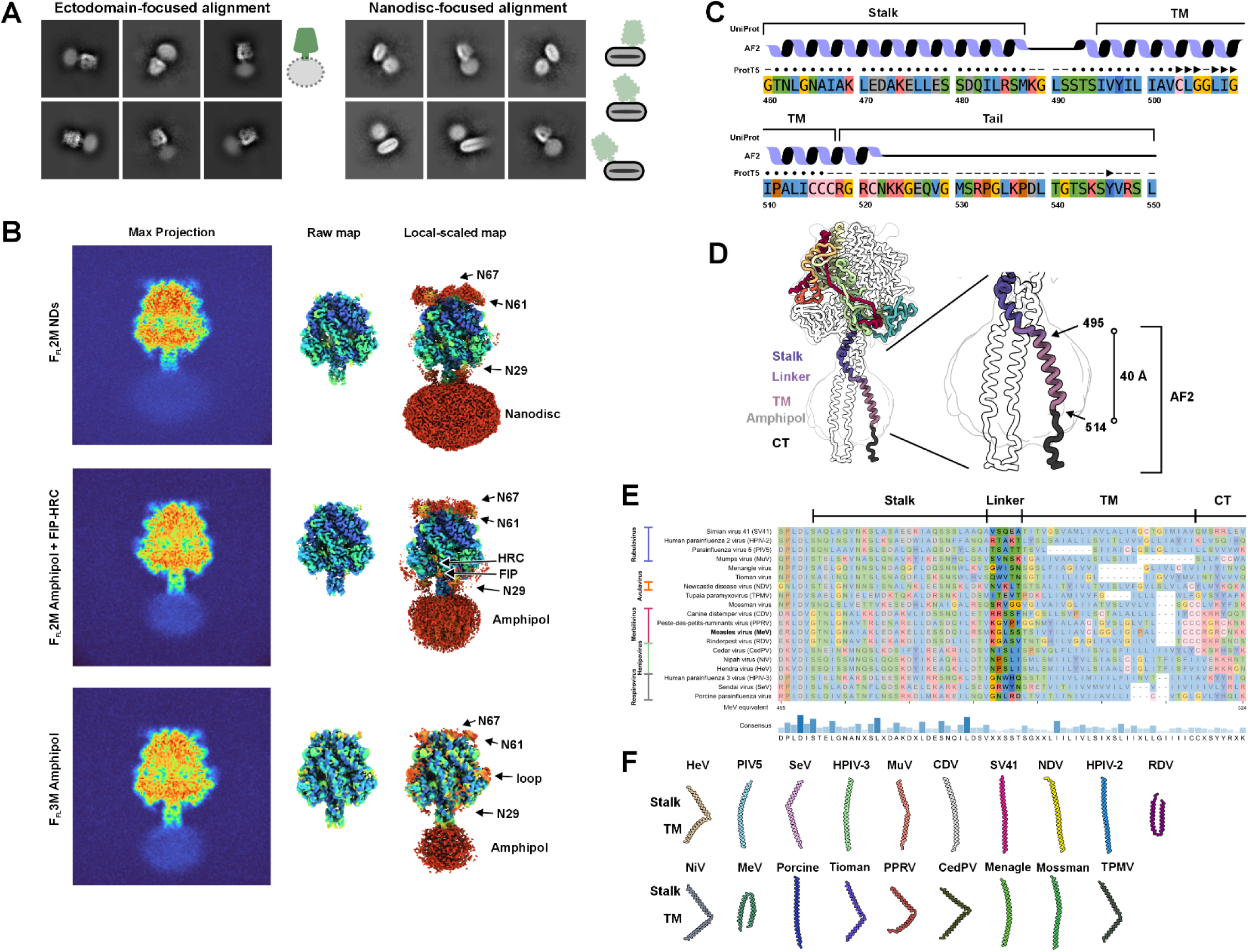
Structural and computational investigation of MeV F transmembrane region. A) Cryo-EM 2D classes of F_FL_ 2M in nanodiscs. Particles were aligned on either the ectodomain (left) or the nanodisc (right). Despite alignment on the nanodisc region, no internal structure (transmembrane region) was observed. The cartoon to the right of the 2D classes illustrates the effect of the alignment on the observed image. In the ectodomain-aligned particles, the nanodisc density is blurred and features are missing. In nanodisc-aligned particles, features inside the nanodisc (lipid bilayer) are visible, but the head domain is blurred and heterogeneously positioned. B) Maximum projection of the final volume (left), raw map (center), and amplified locally scaled map in OccuPy (right) of F_FL_ 2M in nanodisc, F_FL_ 2M + [FIP-HRC]_2_-PEG_11_ in amphipol, and F_FL_ 3M in amphipol, respectively, shows the absence of structural features within the transmembrane region. The amplified locally scaled map, however, reveals clear density for other flexible parts of the structure, such as glycans N29, N61, and N67, as well as partial density for the Δfur loop and flexible parts of [FIP-HRC]_2_-PEG_11_. Local-scale values are shown at the bottom and apply to all maps. C) Computational investigation of the C-terminal part of the F protein. For the F stalk region (residues 460-487), linker (residues 488-492), transmembrane region (TM, residues 495-515), and cytoplasmic tail (CT, residues 516-550), secondary structure was predicted using AlphaFold 2 (AF2) and the protein large language model ProtT5. Both models indicate that the linker region lacks defined secondary structure, similar to the CT. Secondary structure for ProtT5 is depicted as follows: ●: helix, ►: beta sheet, ⁃: loop. D) Hypothetical hybrid model of F_FL_ incorporating the cryo-EM structure of F_FL_ 2M in amphipol (residues 24-485) and AF2 prediction (residues 487-528). The linker was modeled based on the partial density present in the cryo-EM map, and the TM and CT regions were based on AF2 prediction. The conformation of the linker may result in the separation of the TM helices and increase the flexibility of the head region. Additionally, the predicted TM helix spans ∼40 Å, matching the width of the lipid bilayer. E) Multiple sequence alignment (MSA) of representative members of paramyxoviruses. The stalk and TM regions are mapped based on the MeV F sequence. A similar pattern of residues is observed in the linker region across other family members. F) F2 predictions of the stalk-linker-TM region based on the MSA of MeV F. Out of 19 species, more than half show a flexible linker depicted as a kink in the secondary structure. Only one representative structure is shown out of five predicted; some structures displayed mixed straight-bent predictions, but only the bent form is presented.

**Figure S8.**
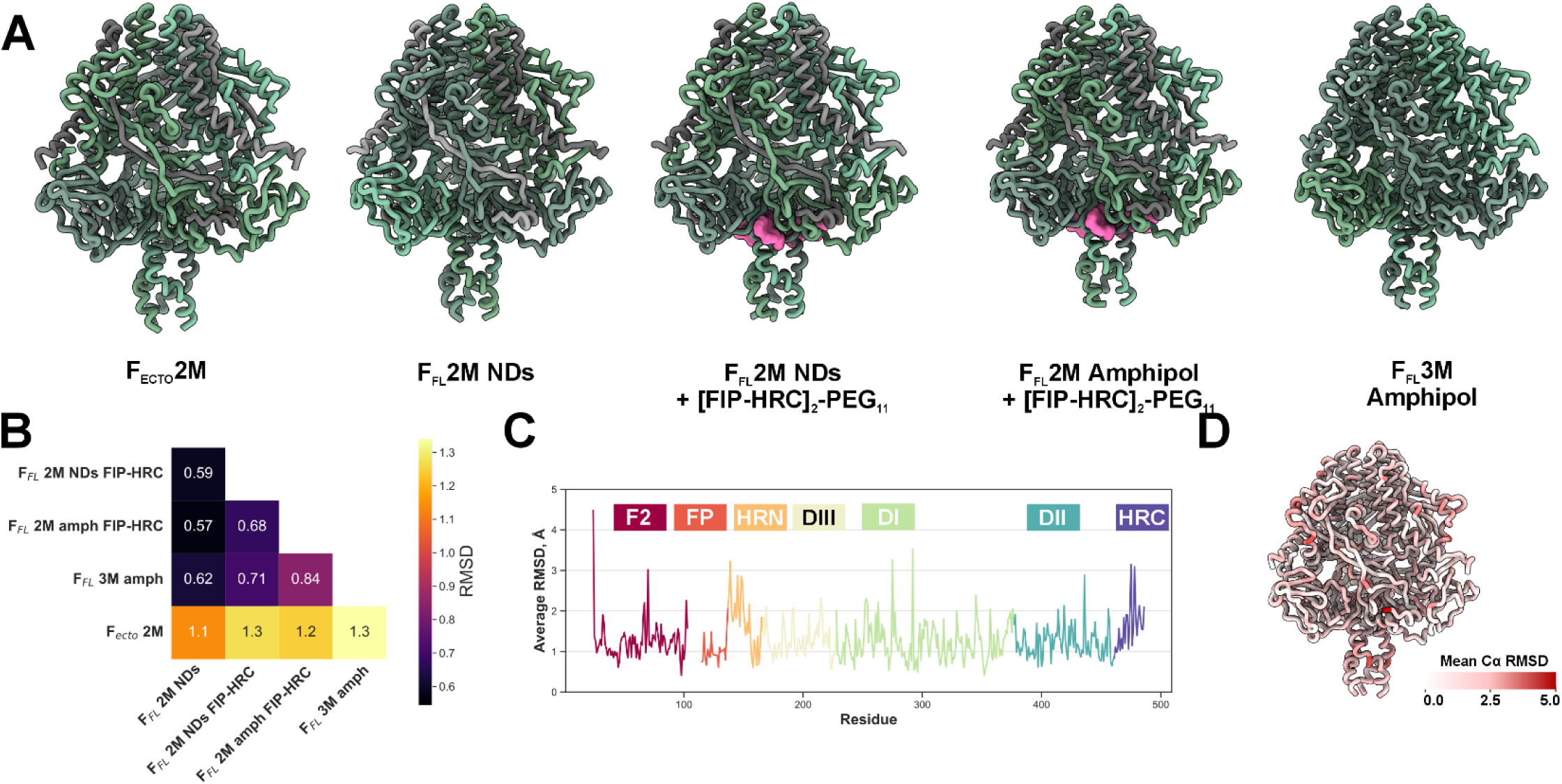
Comparative analysis of F constructs. A) Ribbon representation of F constructs resolved in this manuscript alongside the previously determined F_ECTO_ 2M. The F_2_ chain (2M constructs) is depicted in gray, while the F_1_ and F_0_ chains are illustrated in shades of green. [FIP-HRC]_2_-PEG_11_ density is highlighted in pink. B) Mean Cα RMSD similarity matrix for all structures shown in (A). The color gradient indicates the mean Cα RMSD values. C) Per-residue Cα RMSD plot comparing F_ECTO_ 2M and F_FL_ 2M in nanodisc, indicating regions with local structural differences. The line plot is color-coded by domain. D) Visualization of per-residue Cα RMSD changes from (C) mapped onto the 3D structure of F. The intensity of red color represents the magnitude of the differences.

**Figure S9.**
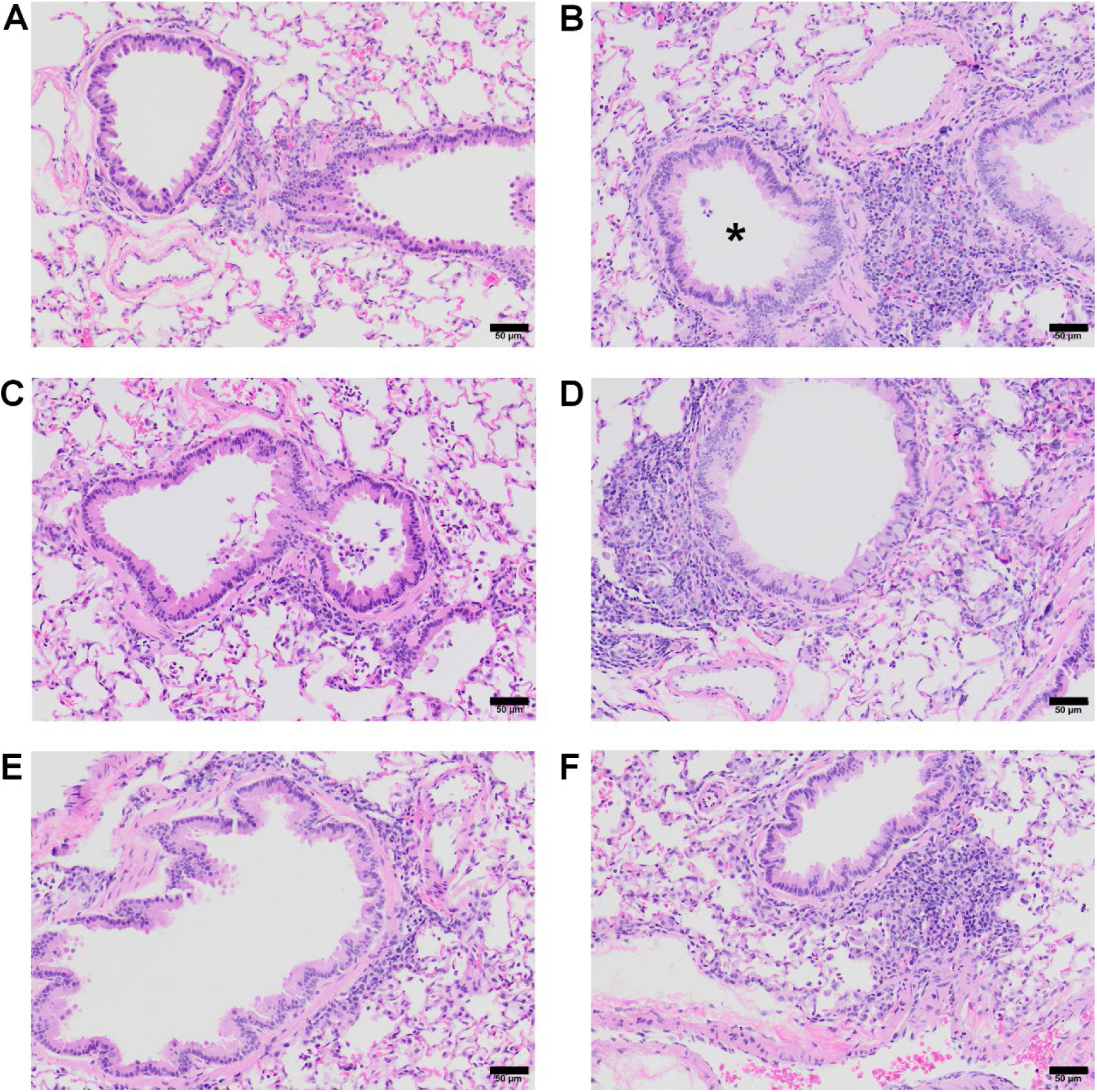
F_FL_ construct immunization does not result in vaccine-enhanced respiratory disease in cotton rats. F protein constructs (5 µg), vehicle control (1 mL 1:1 PBS: AdjuPhos), or measles Edmonston-Zagreb strain (10^4^ TCID_50_) was administered subcutaneously to cotton rats. At day 28 post-primary immunization, F protein constructs or vehicle control were administered again. At day 56 post-primary immunization, cotton rats were infected with the measles virus (wild-type B3-eGFP). Cotton rats were euthanized on day 4 post-infection, and the right lung lobes were prepared for histopathological evaluation. A-F) Representative regions of peribronchiolar inflammatory infiltrates for each treatment or control group. The black bar at the bottom represents a scale of 50 µm. A) The vehicle control group exhibits minimal peribronchiolar infiltrates. B) Edmonston-Zagreb vaccinated animals exhibit more pronounced peribronchiolar inflammatory infiltrates, occasionally forming complete cuffs (*) around affected airways, with a higher proportion of lymphocytes and plasma cells. (C-E) Animals vaccinated and boosted with F protein constructs demonstrate similar or lower degree of peribronchiolar infiltrates compared to Edmonston-Zagreb vaccinated animals (C = F_FL_ 2M; D = F_FL_ 3M). Substantial interstitial and perivascular inflammatory infiltrates are absent.

## Tables

**Table S1.**
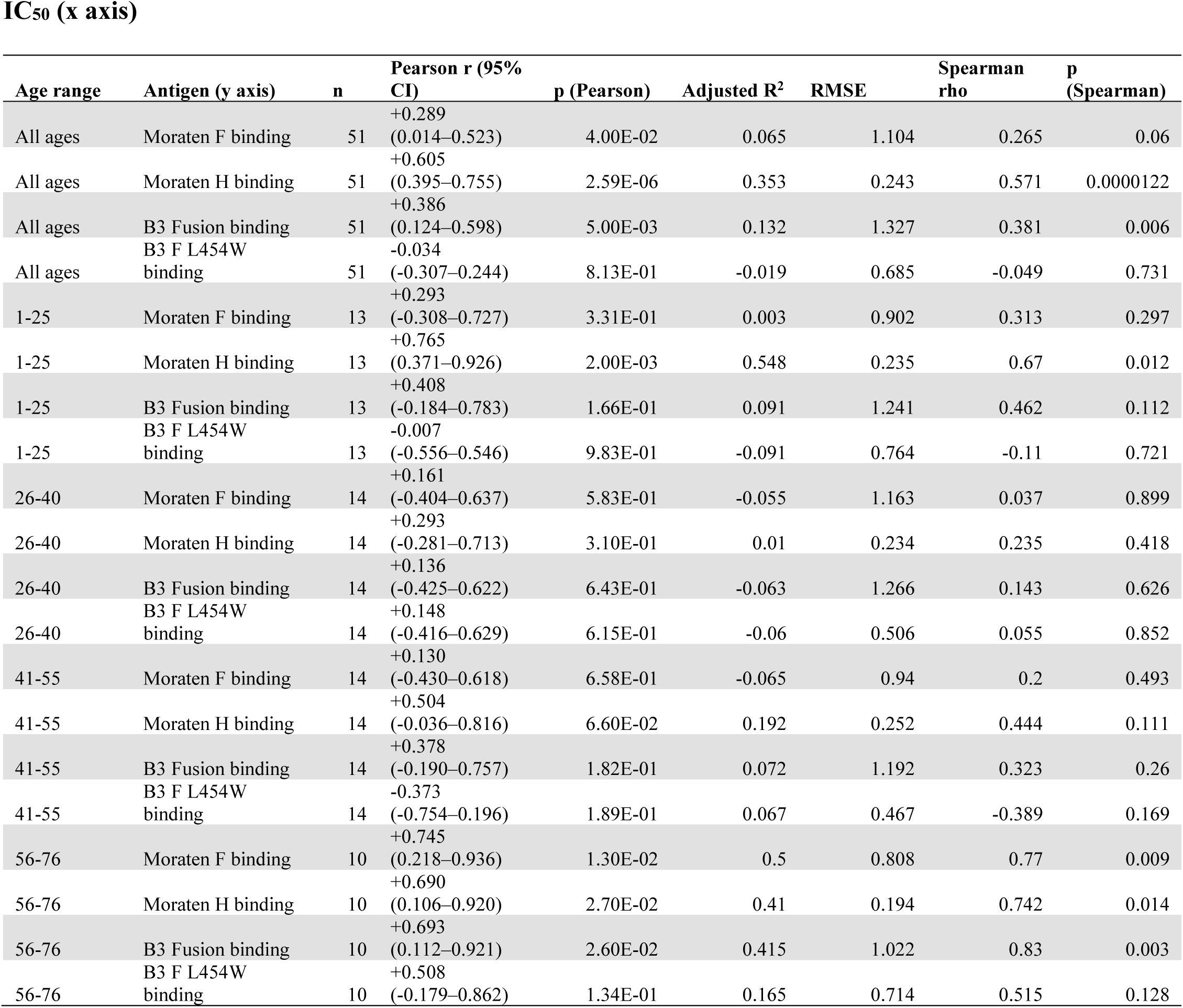
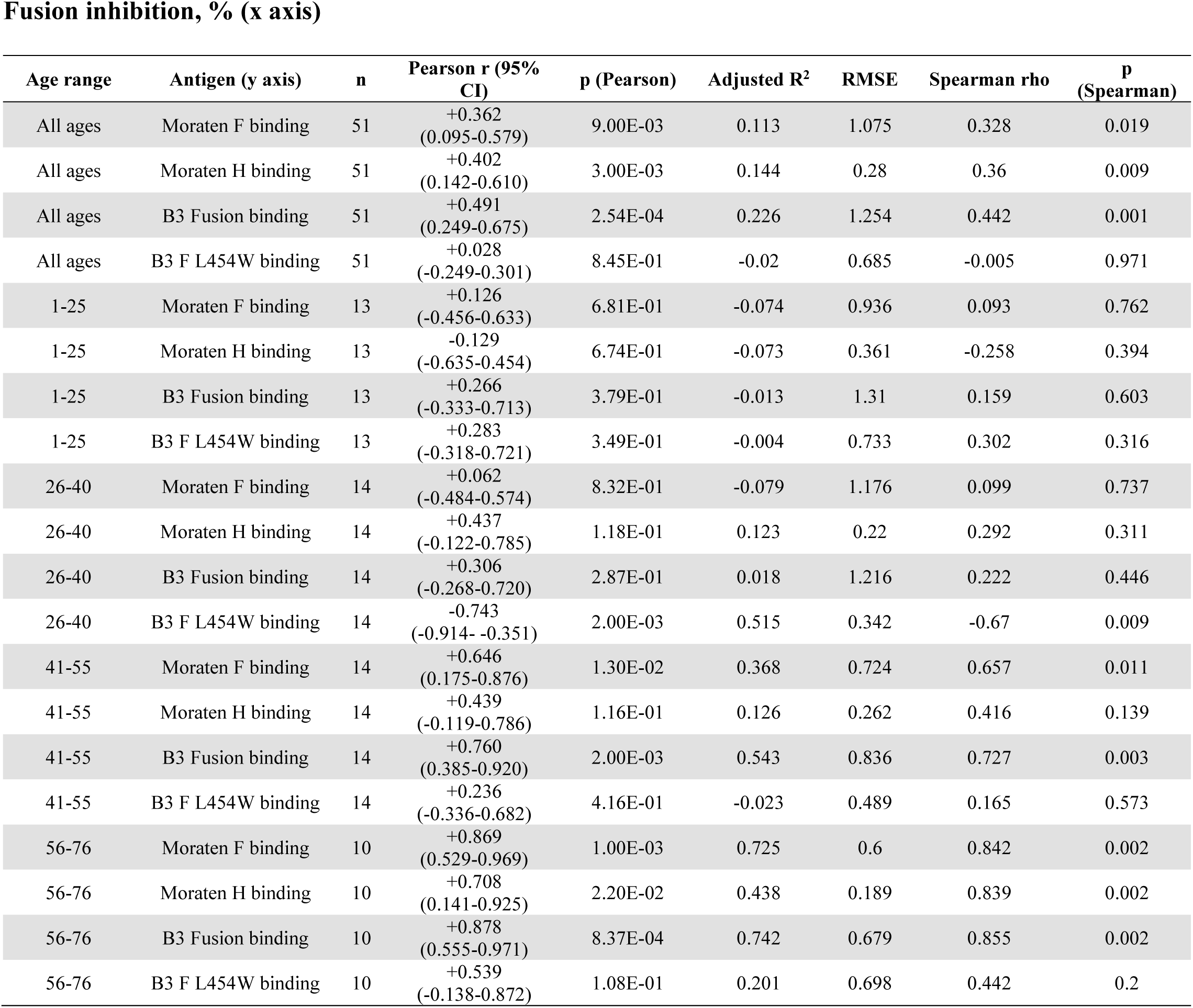
Summary of associations between serum antigen recognition and B3 virus neutralization titers (IC_50_) and fusion inhibition.

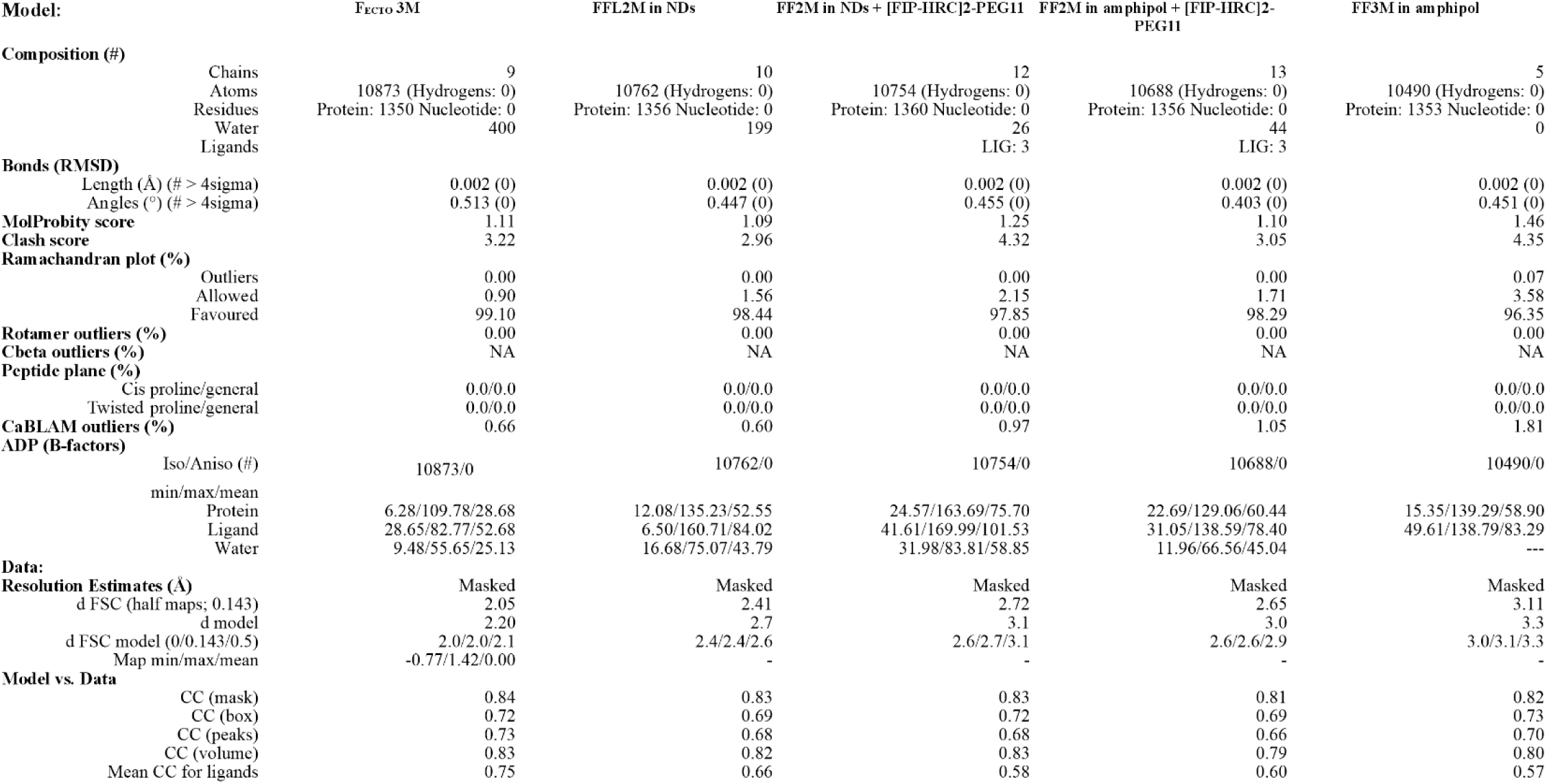
Table S2.

**Table S3:** Semiquantitative histopathologic scoring of inflammatory lesions following measles challenge for severity (0-3) and quantity (0-4) of inflammatory infiltrates.

| Treatment Group;<br>mean (standard<br>deviation) | Peribronchiolar<br>Infiltrates |  | Perivascular<br>Infiltrates |  | Interstitial<br>Infiltrates | Alveolar<br>Infiltrates |
| --- | --- | --- | --- | --- | --- | --- |
|  | Quantity | Severity | Quantity | Severity | Quantity | Quantity |
| Vehicle | 0.75<br>(0.5) | 0.75<br>(0.5) | 0 (0) | 0 (0) | 0 (0) | 0 (0) |
| Edmonston-Zagreb | 2 (1) | 1.4<br>(0.55) | 1.2<br>(0.45) | 1.2<br>(0.45) | 0.2 (0.45) | 0.2 (0.45) |
| F<sub>FL</sub> 2M | 1.8<br>(0.84) | 1.2<br>(0.45) | 0.8<br>(0.84) | 0.8<br>(0.84) | 0.4 (0.55) | 0.6 (0.55) |
| F<sub>FL</sub> 3M | 1.2<br>(0.45) | 1.2<br>(0.45) | 0.8<br>(0.45) | 0.8<br>(0.45) | 0 (0) | 0 (0) |

## Methods

### Clinical sera

Sera from individuals positive for measles antibodies (51 samples, aged 1 to 76, with 18 females and 33 males) were obtained from the Columbia University Biobank (CUB). CUB is an institutional resource that collects, processes, stores, and disseminates biological specimens, biomarkers, and health data for use in biomedical research. Participants in the CUB have consented to allow access to their EHR and will allow re-contact for additional research studies, questionnaires, or collection of additional samples, as well as sharing of their coded data with other academic centers, the NIH, and commercial partners. Researchers who use the CUB data and/or samples are required to adhere to state and local regulations regarding human subject research. The centralized nature of the biobank ensures uniform specimen handling, data quality, and regulatory oversight for biobanking operations. Biological specimens, including blood, urine, and buccal swabs, are processed in the Biobank Resource for Investigating Disease, Genes, and Environment (BRIDGE) laboratory following well-established standard operating procedures and strict quality control criteria.

### Plasmids

The genes of measles morbillivirus (MeV) strains IC323, B3, D8, and vaccine hemagglutinin proteins (H), fusion proteins (F), human nectin 4, and human CD150 proteins were codon-optimized, synthesized, and subcloned into the mammalian expression vector pCAGGS (Epoch Lifescience). A codon-optimized fusion protein sequence derived from the measles virus strain Ichinose-B95a (Taxonomy ID: 645098, Uniprot ID: Q786F3) was synthesized by GenScript and subcloned into the pcDNA3.1 vector. The FL expression constructs were generated by cloning the F open reading frame into a pCG plasmid (Addgene ID #84817^11^) and adding a C-terminal twin Strep-Tag II using the NEBuilder HiFi DNA Assembly. Stabilizing point mutations previously identified in SSPE and MIBE measles strains, including E170G and E455G and Δfur, were introduced sequentially using the Q5 Site-Directed Mutagenesis kit. The Δfur mutation changes the furin cleavage site from RHKR to NHNR, as previously described^58^.

### Cells

HEK293T (human kidney epithelial), Vero (African green monkey kidney), and Vero-SLAM/CD150 cells were grown in Dulbecco’s modified Eagle’s medium (DMEM; Life Technologies; Thermo Fisher Scientific) supplemented with 10% fetal bovine serum (FBS, Life Technologies; Thermo Fisher Scientific) and antibiotics at 37°C in 5% CO_2_. The Vero with human SLAM/CD150 (Vero-hCD150) culture media was supplemented with 1 mg/mL Geneticin (Thermo Fisher Scientific).

HEK 293F cells were cultured in a humidified incubator at 37°C with a 5% CO₂ atmosphere in FreeStyle™ 293 Expression Medium (Gibco). Cells were passaged every 3 to 4 days to maintain optimal cell density and then split to 0.5 × 10^6^ cells/mL in 30 mL of medium. Cells were maintained and used up to the 25^th^ passage.

### Protein expression and purification of F_ECTO_

F_ECTO_ 2M was purified as described previously^35^. F_ECTO_ 3M were expressed in ExpiCHO cells (Thermo Fisher) using the manufacturer’s transfection and high-yield expression protocol. Once cell viability dropped below 60%, typically around day 5, the cells were centrifuged at 4000×g. The medium supernatant was treated with BioLock (IBA Lifesciences) following the manufacturer’s protocol for ExpiCHO medium. After 30 minutes of incubation at room temperature, the medium was centrifuged again at 4000×g and applied to a 5 mL StrepTactin column (Cytiva Life Sciences), pre-equilibrated with HBS buffer (50 mM HEPES, pH 8.0; 150 mM NaCl, pH 8.0), connected to an ÄKTA Pure system (Cytiva Life Sciences). The column was washed with at least 10 column volumes (CV) of buffer or until the absorbance at 280 nm remained constant. Elution was performed using HBS with 5 mM d-desthiobiotin (Sigma). Protein-containing fractions were pooled, concentrated, and flash-frozen in liquid nitrogen for further analysis.

### Protein expression and purification of F_FL_

HEK293F cells were expanded to a volume of at least 500 mL and a density of approximately 1 x 10^6^ cells/mL on the day of transfection. Transfection was performed using the pCG-F_FL_ variant construct at a concentration of 1 μg/mL, using polyethylenimine (PEI, Polysciences) at a 3-fold excess relative to the DNA amount, following a standard protocol. The cells were then cultured for 5 days, and on the morning of the fifth day, they were centrifuged at 4000 × g in 500 mL centrifuge tubes. The resulting cell pellet was resuspended in a volume equivalent to one-tenth of the initial culture (50 mL) in ice-cold HBSE buffer (50 mM HEPES, pH 8.0; 150 mM NaCl; 1 mM EDTA). To this suspension, 200 μL of protease inhibitor cocktail (PIC), 200 μL of BioLock, and 1.5 μL of Benzonase were added, and the mixture was kept on ice. The cell suspension was then transferred to a 15 mL douncer and homogenized by gently moving the piston up and down at least ten times. The homogenized cells were collected into a 50 mL Falcon tube, to which NP-40 (Thermo Fisher) was added to achieve a final concentration of 2%. The cells were incubated at 4°C for 2 hours with gentle rocking at approximately 1 Hz. The solubilized solution was then transferred into 30 mL tubes, and the insoluble fraction was pelleted by centrifugation at 38,000 × g using the SA 20.1 rotor at 4°C for 30 minutes. The supernatant was collected and incubated overnight with 1 mL of StrepTactin resin (Qiagen), previously equilibrated with HBSE, at 4°C with rocking. This mixture was then diluted to 50 mL with HBSE to reduce the final NP-40 concentration. The following day, the mixture was applied to a gravity Bio-Rad column and the resin was washed with 10 mL of HBSE containing 0.06% NP-40 to remove excess NP-40 and wash out contaminants. For nanodisc assembly, protein was eluted in HBSE containing 0.06% NP-40 and 5 mM d-desthiobiotin (Sigma). Alternatively, for amphipol embedding, 1 mL of HBSE containing 0.06% NP-40 was added, and 30 μL of 100 mg/mL Amphipol (Anatrace) was added, and the resin slurry was mixed to allow uniform mixing of the amphipol. The mixture was incubated for 2 hours at 4°C with rocking. Once incubation was finished, resin was washed 3 times with 2 mL of HBSE, and protein was eluted with HBSE with 5 mM d-desthiobiotin. The expression yields of FL constructs varied, ranging from 0.2 mg to 0.8 mg per liter of cell culture.

### MSPD1 purification

MSP1D1 purification was performed based on a previously reported protocol^59^. Briefly, the pMSP1D1 plasmid was obtained from Addgene^60^ was used to heat-shock transform *E. coli* BL21 (DE3) cells (NEB). The cells were then cultured in 1 mL of LB at 37°C with shaking at 200 rpm for 1 hour. The transformed cells were subsequently plated on kanamycin-containing agar plates (50 μg/mL) (ThermoFisher) and incubated overnight at 37°C. A single colony from this plate was used to inoculate 50 mL of LB with kanamycin (50 μg/mL), which was then incubated overnight at 37°C. This starter culture was added to 2 L Terrific Broth (TB) containing 50 μg/mL kanamycin. The culture was grown at 37°C until an A_600_ of 2.5 was reached, induced with 1 mM IPTG, and then the temperature was reduced to 28°C 1 hour post-induction. Cells were harvested 4 hours after induction. For protein purification, the cell pellet was resuspended in lysis buffer (50 mL, 20 mM PBS pH 7.4 with 1% Triton X-100 added post-resuspension) and left on ice for 30 minutes. Lysis was achieved using the M-110P pressure cell disruptor (Microfluidics) at 18k PSI at 4°C. The lysate was then centrifuged (37,000 x g, 20 min) and the supernatant applied to a pre-equilibrated 5 mL Ni-NTA resin (Qiagen) at 4°C. The column was washed with 40 mM Tris, 0.3 M NaCl, 1% Triton X-100 (Anatrace), pH 8.0; followed by 40 mM Tris, 0.3 M NaCl, 50 mM Na-Cholate, 20 mM Imidazole, pH 8.0; and then 40 mM Tris, 0.3 M NaCl, 50 mM Imidazole, pH 8.0. Elution was performed using a buffer containing 40 mM Tris, 0.3 M NaCl, 0.4 M Imidazole, 40 mM Na-Cholate, pH 8.0. Protein purity was validated by SDS-PAGE, and concentration was determined using UV spectroscopy at 280 nm, based on a theoretical extinction coefficient of 21,430 M^-1^ cm^-1^.

### Nanodisc assembly

The concentration of the eluted F_FL_ constructs in HBSE buffer supplemented with 0.06% NP-40 and 5 mM D-desthiobiotin was determined by measuring the absorbance at 280 nm. Typical concentration values were around 0.1 mg/mL. Due to potential inaccuracies in protein concentration measurements in the presence of NP-40, nanodisc assembly was conducted with a 5-fold molar excess of nanodiscs relative to the protein trimer. Lipids, specifically POPC and POPG (Avanti), were mixed in an 80:20 molar ratio, extracted from chloroform solutions, and transferred to a glass test tube. The lipids were then dried under an argon stream for 15 minutes. To ensure complete removal of chloroform, the dried lipids were further washed with 1 mL diethyl ether and dried for an additional 10 minutes. Then, the lipids were resuspended in 30 mM DDM (Anatrace) in water, followed by vigorous vortexing until they were fully solubilized. This solution underwent two freeze-thaw cycles, alternating between liquid nitrogen and 37°C, and was then sonicated for 15 minutes in a water bath to ensure a homogenous, clear lipid suspension. The lipid mixture was added to the protein solution at a 300-fold molar excess relative to the protein, and the mixture was incubated for 15 minutes. Subsequently, a 5-fold molar excess of MSPD1 relative to the protein was added, followed by another 15-minute incubation. To remove NP-40, 10% of the total volume of wet Biobeads SM2 (Bio-Rad) was added, and the mixture was incubated overnight with rocking at 1 Hz. The Biobeads were replaced the following day until no change in absorbance at 280 nm was detected, indicating the successful removal of NP-40. Empty nanodiscs were separated from protein-embedded nanodiscs through size exclusion chromatography using a Superdex 200 10/300 GL column (GE Healthcare). The successful incorporation of the protein into the nanodiscs was verified using SDS-PAGE gel electrophoresis and NS EM.

### Protein concentration determination

Protein concentration was determined using absorbance measurements at 280 nm using a Nanodrop 2000. The spectrophotometer was blanked on water, and the buffer correction was done externally. For molar concentrations, given extinction coefficients were used: F_FL_ (all variants): 51,800 cm^-1^ M^-1^, F_ECTO_ (all variants): 48,820 cm^-1^ M^-1^.

### Stability measurements using nano-differential scanning fluorimetry (nanoDSF) and dynamic light scattering (DLS)

Stability measurements, expressed as apparent melting temperature and changes in hydrodynamic radius as a function of temperature, were conducted using the Nanotemper Prometheus Panta. Three 10 μL capillaries, each filled with 0.2 mg/mL of protein for each construct, were subjected to a temperature ramp of 1°C/min from 25°C to 95°C. Protein unfolding was monitored through changes in the nanoDSF fluorescent signal, with the first derivative used to determine the apparent melting temperature. DLS signals were simultaneously recorded. Data analysis was performed using PR.Panta 1.7.4 software (Nanotemper), and the averages of processed data, including the first derivative of the 330 nm signal and the cumulant fit of the DLS data, were plotted using Python.

### Grid preparation

The freezing procedure and grid types were consistent for all protein constructs, following the protocol outlined previously^35^. Briefly, Quantifoil R2/2 grids were washed with ethyl acetate the day before freezing. On the day of freezing, graphene oxide (GO, Sigma) was freshly prepared and applied to the grid. Specifically, 3 μL of 0.2 mg/mL GO was applied and incubated for 3 minutes, followed by blotting and washing with 20 μL ddH2O. Afterward, the grid was left to dry for 10 minutes. Within the Vitrobot Mark IV, protein samples at a concentration of 0.15-0.25 mg/mL were applied to the grid at 4°C and 100% humidity, incubated for 15 seconds, and then blotting was performed for 3 seconds before the sample was flash-frozen in liquid ethane cooled with liquid nitrogen.

### Structure determination F_ECTO_ 3M

All samples were collected using a Titan Krios microscope operating at 300kV with a Gatan K3 camera and energy filter set to 20 eV, resulting in a magnification of 130,000X and a pixel size of 0.66 Å/px. The exposure settings were 70 ē/Å^2^ with an exposure rate of 15 ē/px/s. All data processing was carried out using cryoSPARC version 4.5. A total of 10,974 TIFF movies were recorded in super-resolution mode with 2x binning using the EPU software. These movies underwent motion correction, and contrast transfer function (CTF) parameters were estimated using the Patch CTF algorithm. Initially, particle picking was performed using a blob picker with a radius range of 80-120 Å. Particles were extracted at a box size of 512 px, Fourier-cropped to 64 px, and subjected to 2D classification using the Deep2D classification protocol^35^. After selecting well-defined 2D classes, a subset of the 10,000 best particles was used to train the Topaz model^61^. The trained Topaz model was then used for particle picking, followed by another round of Deep2D classification to remove suboptimal particles. An initial model was generated using the Ab Initio protocol. Subsequently, all selected particles from the 2D classification were aligned to this volume using Homogeneous and Non-Uniform Refinement (NUR) protocols, followed by 3D classification to exclude suboptimal particles. The best 3D classes were further refined using NUR. When the Nyquist resolution imposed by the box size was reached, the particles were unbinned to 128 px, and the same refinement and 3D classification protocols, including NUR, were repeated until the resolution limit was reached. Finally, particles were extracted at 512 px, Fourier-cropped to 384 px, and further underwent CTF refinement, reference-based motion correction, and additional 3D classifications to discard remaining suboptimal particles. The selected particles underwent symmetry expansion and were refined using the local refinement protocol, followed by another round of 3D classification. The best class was ultimately refined using local refinement, resulting in a final resolution of FSC_0.143_ = 2.05 Å.

### Structure determination F_FL_

The exposure settings for F_FL_ datasets were 50 ē/Å^2^ with an exposure rate of 15 ē/px/s. A summary of the data collection statistics can be found in **Table S2**. The approach described in^35^ was followed, and recorded movies in TIFF format were motion corrected using Relion 4’s own algorithm. CTF values were estimated using CTFfind 4. Initial particle picking was performed using the LoG picker inside Relion and followed by 2D classification and Topaz picking. Particles were extracted with binning to 64 pixels, and 2D classification was carried out in cryoSPARC. Particles were then moved back to Relion for initial model generation and 3D classification. The 3D classification was continued until class equilibrium was reached, as monitored by the Follow Relion Gracefully program^62^. Only classes with expected features were selected, and once the Nyquist resolution in the binned data was achieved, particles were unbinned to 128 pixels for another round of 3D classification. The selected 3D class was run in a 3D refinement protocol, and the final particle set was unbinned to ∼3/4 of the original box size. Further processing involved Refine 3D, CTFrefine, and Bayesian Polishing in Relion. Finally, the final particle stacks were imported to cryoSPARC, where Non-Uniform Refinement, symmetry expansion to C3, and Local Refinement were used.

### Glycoproteomics LC-MS/MS measurements

The experiments were performed as previously described^35,63^ with minor modifications, as follows. Recombinant measles F was denatured in buffer with 2% SDS, 200 mM Tris/HCl, 10 mM tris(2-carboxyethyl)phosphine, pH 8.0, at 95°C for 10 minutes, followed with a 30 minute reduction at 37°C. Samples were next alkylated by adding 40 mM iodoacetamide and incubated in the dark at room temperature for 45 minutes. Three micrograms of recombinant F was used for each protease digestion. Samples were split in half for parallel digestion with chymotrypsin (Sigma) and alpha lytic protease (Sigma). Sera-Mag beads (GE Healthcare) were used at a bead/protein ratio of 10:1 (w/w) for protein binding. Ethanol was added to the protein-beads mixture reaching final concentration 80% to enhance binding. The mixture was incubated at RT for 10 minutes at 1000 rpm in a ThermoMixer. The supernatant was removed, and protein-bound beads were washed 3 times with 80% ethanol. MeV F was digested on beads overnight at 37°C in 25 μL 50 mM ammonium bicarbonate at 1000 rpm in a ThermoMixer. The supernatant was taken for MS analysis. For each sample and protease digestion, approximately 50 ng of peptides were run by online reversed phase chromatography on a Dionex UltiMate 3000 (Thermo Fisher Scientific) coupled to a Thermo Scientific Orbitrap Fusion mass spectrometer. A Poroshell 120 EC C18 (50 cm × 75 µm, 2.7 µm, Agilent Technologies) analytical column and a ReproSil-Pur C18 (2 cm × 100 µm, 3 µm, Dr. Maisch) trap column, were used for peptide separation. Duplicate samples were analyzed with two different mass spectrometry methods, using identical LC-MS parameters and distinct fragmentation schemes. In one method, peptides were subjected to Electron Transfer/Higher-Energy Collision Dissociation (HCD) fragmentation. In the other method, all precursors were subjected to HCD fragmentation, with additional EThcD fragmentation triggered by the presence of glycan reporter oxonium ions. A 90-minute LC gradient from 0% to 44% acetonitrile was used to separate peptides at a flow rate of 300 nL/min. Data was acquired in data-dependent mode. Orbitrap Fusion parameters for the full scan MS spectra were as follows: a standard AGC target at 60,000 resolution, scan range 350–2000 m/z, Orbitrap maximum injection time 50 ms. The ten most intense ions (2+ to 8+ ions) were subjected to fragmentation. For the EThcD fragmentation scheme, the supplemental higher energy collision dissociation energy was set at 27%. MS2 spectra were acquired at a resolution of 30,000 with an AGC target of 800%, maximum injection time 250 ms, scan range 120–4000 m/z and dynamic exclusion of 16 s. For the triggered HCD-EThcD method, the LC gradient and MS1 scan parameters were identical. The ten most intense ions (2+ to 8+) were subjected to HCD fragmentation with 30% normalized collision energy from 120–4000 m/z at 30,000 resolution with an AGC target of 100% and a dynamic exclusion window of 16 s. Scans containing any of the following oxonium ions within 20 ppm were followed up with additional EThcD fragmentation with 27% supplemental HCD fragmentation. The triggering reporter ions were: Hex(1) (129.039; 145.0495; 163.0601), PHex(1) (243.0264; 405.0793), HexNAc(1) (138.055; 168.0655; 186.0761), Neu5Ac(1) (274.0921; 292.1027), Hex(1)HexNAc(1) (366.1395), HexNAc(2) (407.166), dHex(1)Hex(1)HexNAc(1) (512.1974), and Hex(1)HexNAc(1)Neu5Ac(1) (657.2349). EThcD spectra were acquired at a resolution of 30,000 with a normalized AGC target of 400%, maximum injection time 250 ms, and scan range 120–4000 m/z.

### Bioinformatics analysis of membrane membrane-proximal region

Secondary structure predictions for the stalk, membrane-proximal (linker), and transmembrane regions were performed using AlphaFold2 via ColabFold^64^ and ProtT5^65^. Representative sequences from the Paramyxoviridae family were obtained from the NCBI Protein database using the search query: “((“Paramyxoviridae”[Organism] OR paramyxoviridae[All Fields]) AND fusion[All Fields]) AND swissprot[filter]”, selecting one sequence per virus. Sequences were aligned using Clustal Omega^66^, with the measles F protein (Q786F3) chosen as the reference. The regions corresponding to the stalk, linker, and transmembrane domains were selected for all representative sequences and processed using the ColabFold monomer batch mode, generating 15 structure predictions per sequence (3 independent runs). Structure selection was based on the presence of a helix break in at least one prediction, classifying the sequence as having a helix discontinuity.

### Glycoproteomics data analysis

The acquired data was analyzed using Byonic (v4.5.2) against a custom database of recombinant F and the proteases used in the experiment, searching for glycan modifications with 12/24 ppm search windows for MS1/MS2, respectively. Up to ten missed cleavages were permitted using C-terminal cleavage at F/Y/W/M/L for chymotrypsin, or T/A/S/V for alpha lytic protease. For N-linked analysis, carbamidomethylation of cysteine was set as fixed modification, oxidation of methionine/tryptophan as variable common 1, and hexose on tryptophan as variable rare 1. N-glycan modifications were set as variable common 2, allowing up to maximum 2 variable common and 1 rare modification per peptide. All N-linked glycan databases from Byonic were merged into a single non-redundant list to be included in the database search. All reported glycopeptides in the Byonic result files were manually inspected for quality of fragment assignments (with scores ≥ 200). All glycopeptide identifications from both EThcD and HCDpdEThcD runs were merged into a single non-redundant list per sequon. Glycans were classified based on HexNAc content as truncated (≤ 2 HexNAc; < 3 Hex), paucimannose (2 HexNAc, 3 Hex), oligomannose (2 HexNAc; > 3 Hex), hybrid (3 HexNAc) or complex (> 3 HexNAc). Byonic search results were exported to mzIdentML format to build a spectral library in Skyline (v22.2.0.312) and extract peak areas for individual glycoforms from MS1 scans. The full database of variable N-linked glycan modifications from Byonic was manually added to the Skyline project file in XML format. Reported peak areas were pooled based on the number of HexNAc, Fuc or NeuAc residues to distinguish truncated, paucimannose, oligomannose, hybrid, and complex glycosylation, or the degree of fucosylation and sialylation, respectively.

### Glycan shielding

Glycan shielding was assessed using the GlycoSHIELD program^67^, with the F_FL_ 2M structure serving as the template. For each glycosylation site, the nearest related glycan was assigned: N29 with high mannose, N61 with complex glycan lacking fucose, and N67 with complex glycan containing fucose. The combined structure was used to calculate solvent accessibility using the provided scripts, and figures were generated using ChimeraX.

### Statistical analysis

Data analysis and visualizations were conducted using a Python notebook integrating SciPy, Matplotlib, Pandas, Seaborn, and Statannotations package^68^. Statistical significance was determined using the Mann-Whitney test. For p-value annotations, the following symbols were used: ‘ns’ (not significant) for a corrected p-value (p-adj) > 0.05, * for 0.01 < p-adj ≤ 0.05, ** for 0.001 < p-adj ≤ 0.01, *** for 0.0001 < p-adj ≤ 0.001, and **** for p-adj ≤ 0.0001.

### Cell surface staining with conformation-specific anti-F mAbs and sera from in vivo immunization

*Evaluating F thermal stability:* HEK293T cells were transiently transfected with viral glycoprotein constructs: B3 wild-type F(GS63325-1), D8 wild-type F (GS77420-3), B3 L454W F (GS55608), B3 E455G F (GS74446-1), Moraten F (GS55588-1, alias vaccine F), and Moraten H (GS55588-2, alias vaccine H), Edmonston-Zagreb F (GS77870-4), AIK-C F (GS77870-1), CAM-70 F (GS77870-2), Shanghai-191 F (GS77870-5) and Changchun-47 F (GS77870-3), and kept at 37 °C for 18-24 hours. For thermal stability, the cells expressing the F proteins were incubated at the indicated temperatures and times. Cells were then incubated with F-specific mouse monoclonal antibodies targeting either prefusion (mAb77^35^) or post-triggered (mAbC6^38^) conformations of F for 1 hour on ice, then washed with PBS and incubated for 1 hour on ice with Alexa Fluor 488 anti-mouse secondary antibodies (Life Technologies, dilution 1:1,000). Cells were washed with PBS and fixed for 10 minutes on ice with 4% paraformaldehyde (PFA) with a 1:1,000 dilution of DAPI (4′,6-diamidino-2-phenylindole; Thermo Fisher) for 1 hour. Plates were imaged using a Cytation 5, and the relative fluorescence of antibody-bound cells was determined with a BioTek Gen5. The total number of positive cells were counted. Thermal stability is represented by the number of positive cells for mAb77 or mAbC6.

### Quantifying levels of prefusion versus post-triggered antibodies in vaccinated CR sera

Sera from CRs immunized with control vehicle (PBS) or the indicated F proteins were diluted 1:50, followed by serial two-fold dilutions. Serially diluted sera were overlaid on HEK293T cells transiently transfected with wild-type F that were either maintained at 37°C or incubated at 55 °C for 30 minutes immediately prior to sera addition (to induce F’s triggering into the postfusion conformation). After sera was incubated with cells for 1 hour on ice, cells were washed with PBS and incubated with protein G Alexa Fluor 488 conjugate (1:1,000; Life Technologies) for 1 hour on ice, washed with PBS, and fixed for 10 minutes on ice with 4% paraformaldehyde with a 1:1,000 dilution of DAPI (4′,6-diamidino-2-phenylindole; Thermo Fisher). Plates were imaged using a Cytation 5, and the numbers of fluorescent antibody-bound cells were quantified with a BioTek Gen5. The graphical representation uses bars to represent means and error bars to indicate the 95% confidence interval.

### Quantifying levels of specific anti-H and F antibodies in human sera

Human sera from the Columbia Biobank (BRIDGE) were diluted 1:50 and overlaid on HEK293T cells transiently transfected with Measles: B3 wild-type F(GS63325-1), B3 L454W F (GS55608), Moraten F (GS55588-1, alias vaccine F), and Moraten H (GS55588-2, alias vaccine H), and were maintained at 37°C for 18-24 hours. After a 1-hour incubation on ice with cell-sera, the cells were washed with CO2-independent media and then incubated with protein G Alexa Fluor 488 conjugate (1:1000; Thermo Fisher) for an additional 1 hour on ice. Then they were washed three times with CO_2_-independent media and fixed for 10 minutes on ice with 4% paraformaldehyde with a 1:1,000 dilution of DAPI (4′,6-diamidino-2-phenylindole; Thermo Fisher). Plates were imaged using a Cytation 5, and the total well fluorescence signals were quantified with a BioTek Gen5 (version 3.16).

### β-Galactosidase (β-Gal) complementation-based fusion assay

The β-Gal complementation-based fusion assay was performed as described in^30,37,41^. Briefly, HEK293T cells were transiently transfected with either hCD150-venus or venus and the β-Gal omega subunit. After 5h these cells were mixed with HEK293T cells co-expressing the β-Gal alpha subunit together with several combinations of viral glycoproteins: (1) measles H and F, (2) chimera of H with Newcastle disease virus (NDV) head domain and measles stalk domain described in^37^ with measles F, and chimera of H with NDV head domain and measles stalk domain with canine distemper virus F (the measles H stalk effectively activates CDV F, and therefore this construct served as a negative control since measles H stalk specific antibodies in the human sera would not block CDV fusion), in the presence or absence of 1:200 dilute human sera. Fusion was stopped by lysing the cells, and after the addition of the substrate (Tropix Galacto-Star chemiluminescent reporter assay system, Applied Biosystems), the cells were incubated for 1 hour and then quantified using either a Tecan M1000 microplate reader for luminescence or a Biotek Cytation 5 for imaging.

### Human serum neutralization assay

Two-fold human sera dilutions with a starting dilution of 1:50 were incubated with 350 PFU/mL of MeV wild-type B3-mCherry virus on Vero hCD150 cells. Three days later, the plates were imaged, and the plaques were counted using a Cytation 5.

### Correlation analysis and plotting

Correlation plots were generated using raw or processed data. Raw data included serum binding values directly measured from plates, while processed data included results from the fusion inhibition and viral neutralization assays. For the fusion inhibition assay, data was normalized against the no-serum control using the equation:

Fusion inhibition (%) = 1−(condition/Not treated)×100.

For the sera viral neutralization assay, each plate was analyzed individually to account for potential plate-specific variability. Data was fitted using a four-parameter logistic function with globally shared minimum and maximum values (A1 and A2) across all sera dilutions on the plate. The resulting EC_50_ values were used for subsequent analyses. Correlation plots were fitted using linear regression, implemented via the *regplot* function in the seaborn library, with statistical parameters calculated using SciPy. Significance was set at a threshold of p < 0.05. Regression lines were displayed with a solid line, and a transparent overlay indicated a 95% confidence interval. Pairwise comparisons were visualized using the seaborn pairplot function.

### Viral rescue

HEK293T cells were transfected using Lipofectamine 2000 on 6-well plates coated with poly-D-lysine. The transfection mixture included purified plasmid constructs coding for viral proteins N (GS6558900-1), P (GS59482), and L (GS58944), all sourced from Epoch Life Science, as per our design specifications. Additionally, purified plasmids for T7 polymerase (GS58929) and the genome for the MeV B3 eGFP wild-type virus (GS68838) or the MeV B3 mCherry wild-type virus (GS69677-10) were included. Cells were incubated overnight at 37°C in Opti-MEM medium. The medium was then replaced with DMEM supplemented with 10% FBS. Cells were subjected to a heat shock treatment by incubating them at 42°C in a water bath for 3 hours. Subsequently, the cells were returned to a 37°C incubator for 48 hours. After this incubation, the HEK293T cells were detached and transferred to Vero-hCD150 cells in T-75 flasks to facilitate the amplification of the rescued virus. The virus underwent four passages and was titrated to a concentration of 10^6^ plaque-forming units per milliliter (PFU/mL). RNA from the virus sample was then sequenced, and the virus stock was used for subsequent *in vivo* or *in vitro* experiments (see ^35,38^).

### Neutralization assay for cotton rat sera (NT)

Two-fold serum dilutions with a starting dilution of 1:10 were incubated with 25 TCID_50_ of MeV wild-type B3-eGFP virus for 1 hour at 37 °C and plated in duplicate onto 104 Vero-hCD150 cells in MEM with 2% newborn calf serum in a 96-well plate. Three days later, viral titers were determined microscopically by cytopathic effect (CPE) and eGFP expression.

### In vivo experiments: cotton rats

The molecularly cloned wild-type MeV (strain B3 wild-type eGFP)^35^ was grown on BJAB cells and titrated on Vero-hCD150 cells using the tissue culture infectious dose 50 method. The vaccine virus strain Edmonston-Zagreb was obtained from the CDC measles virus collection and grown and titered on Vero cells.

Inbred cotton rats (*Sigmodon hispidus*) were purchased from Inotiv, Inc., Indianapolis. Both male and female cotton rats aged 5 to 7 weeks were used. Animals received subcutaneously either 2 or 5 µg of the indicated proteins, 10^4^ TCID_50_ of the MeV live vaccine (Edmonston-Zagreb), or the vehicle. The animals were immunized twice and were challenged two weeks after the last immunization. For intranasal infection, 10^5^ TCID_50_ of MeV (strain B3 wild-type eGFP) was inoculated intranasally to isoflurane-anesthetized cotton rats in a volume of 100 µL. Four days after infection, the animals were euthanized by CO_2_ inhalation. All right lung lobes and accessory lung lobes were collected for histologic evaluation. Left lung lobes were collected, weighed, and homogenized with a glass Dounce homogenizer for viral titration. Serial 10-fold dilutions of supernatant fluids were assessed on Vero-hCD150 cells in 48-well plates for the presence of infectious virus, using CPE as the endpoint. Plates were scored for CPE microscopically after 5 days, and the tissue culture infectious dose 50 (TCID_50_) was calculated. The results are presented as TCID_50_/g of lung tissue.

### Histology preparation and evaluation

Right cranial, right middle, right caudal, and accessory lung lobes were perfused and fixed in 10% neutral buffered formalin. Standardized sections were trimmed and paraffin embedded. 4-µm-thick sections were stained with H&E and graded histologically using a semi-quantitative scoring system for inflammatory infiltrates. The quantity of peribronchiolar and perivascular infiltrates were evaluated based on the proportion of profiles with associated inflammatory infiltrates (0% = grade 0; 0-25% = grade 1; 25-50% = grade 2; 50-75% = grade 3; >75% = grade 4). Severity of peribronchiolar and perivascular infiltrates were evaluated on a severity scale based on the most severely affected profile (no affected profiles = grade 0; interrupted cuff = grade 1; complete cuff <5 cell layers thick = grade 2; complete cuff >5 cell layers thick = grade 3). Interstitial and alveolar infiltrates were graded on the quantity of alveolar septae or alveolar spaces, respectively, with inflammatory infiltrates (0% = grade 0; 0-10% = grade 1; 10-33% = grade 2; 33-66% = grade 3; >66% = grade 4).

## Data availability

Structural models have been deposited in the Protein Data Bank (PDB, https://www.rcsb.org/) and the cryo-EM maps in the Electron Microscopy Data Bank (EMDB, https://www.emdataresource.org/). The available structures include: F_ECTO_ 3M (PDB ID: 9DTU, EMDB ID: EMD-47166); F_FL_ 2M in nanodisc (PDB ID: 9COE, EMDB ID: EMD-45777); F_FL_ 2M bound by [FIP-HRC]2-PEG11 in nanodisc (PDB ID: 9COF, EMDB ID: EMD-45778); F_FL_ 2M bound by [FIP-HRC]2-PEG11 in amphipol (PDB ID: 9COG, EMDB ID: EMD-45779); and F_FL_ 3M in amphipol (PDB ID: 9COH, EMDB ID: EMD-45780).

The raw LC-MS/MS files and glycopeptide identifications have been deposited to the ProteomeXchange Consortium via the PRIDE partner repository with the dataset identifier PXD050366. Post-publication, all research materials referenced in this study will be made available upon reasonable request through Material Transfer Agreements (MTAs).

## Contributions

DSZ: Designed protein constructs; cloned and expressed proteins; recorded all negative stain and cryo-EM data; processed and solved structures; wrote the code for analysis; analyzed all data presented in the manuscript; prepared figures; wrote the manuscript. GZ: Performed virological and cell biology experimentation; analyzed data; contributed to manuscript writing. RDM: Performed virological and cell biology experimentation; analyzed data. DL: Performed cell-based experiments with sera and analyzed the data. GN: Assisted with screening, protein purification, and negative stain electron microscopy. WP: Conducted glycosylation pattern analysis. CP: Purified proteins. KS: Analyzed data. DT: Produced and purified proteins; contributed to manuscript editing. JK: Conducted molecular modeling. MA: Assisted with data curation, analysis, and manuscript writing. LDC: Performed virological and cell biology experimentation; analyzed data. CL: Conducted *in vivo* experimentation and analyzed data. AD: Performed chemical engineering. DP: Assisted with cloning, expression, and purification of constructs. RDA: Assisted with cryo-EM data acquisition; maintained microscopes. JM: Conducted pathology assessment. GM: Conducted molecular modeling. NVD: Supervised the work; contributed to manuscript writing. JS: Conducted glycosylation pattern analysis. CAA: Supervised the work; analyzed data. KMH: Analyzed data; contributed to manuscript writing and figure preparation. AM: Analyzed and interpreted results; contributed to manuscript writing. ALG: data analysis and interpretation, and manuscript writing. SN: Conducted *in vivo* experimentation, data analysis, and interpretation; supervised the work; contributed to manuscript writing. EOS: Analyzed and interpreted results; supervised the work; secured funding; contributed to manuscript writing. MP: Designed proteins and experiments; performed virological and cell biological experimentation; analyzed and interpreted results; supervised the work; secured funding; manuscript writing.

## Ethics declarations

MP, AM, DSZ, and EOS have a provisional patent related to thermostable MeV F proteins as a vaccine. All the authors declare that they have no competing interests in relation to the work described in this manuscript.

